# Single-cell analysis reveals MHCII expressing keratinocytes in pressure ulcers with worse healing outcomes

**DOI:** 10.1101/2021.04.20.440591

**Authors:** Dongqing Li, Shangli Cheng, Yu Pei, Pehr Sommar, Jaanika Kärner, Eva K. Herter, Maria A. Toma, Letian Zhang, Kim Pham, Yuen Ting Cheung, Xingqi Chen, Liv Eidsmo, Qiaolin Deng, Ning Xu Landén

## Abstract

Pressure ulcer (PU) is a chronic wound often seen in spinal cord injury patients and other bed-bound individuals, particularly in the elderly population. Despite its association with high mortality, the pathophysiology of PU remains poorly understood. Here, we compared single-cell transcriptomic profiles of human epidermal cells from PU wound edges with those from uninjured skin and acute wounds (AWs) in healthy donors. We identified significant shifts in the cell composition and gene expression patterns in PU. In particular, we found that major histocompatibility complex class II (MHCII) expressing keratinocytes were enriched in patients with worse healing outcomes. Furthermore, we showed that the IFNγ in PU-derived wound fluid could induce MHCII expression in keratinocytes and that these wound fluid-treated keratinocytes inhibited autologous T cell activation. In line with this observation, we found that T cells from PUs enriched with MHCII+ keratinocytes produced fewer inflammatory cytokines. Overall, our study provides a high-resolution molecular map of human PU compared to AW and intact skin, providing new insights into PU pathology and the future development of tailored wound therapy.

## Introduction

A pressure ulcer (PU) is a chronic nonhealing wound that is caused by the continuous pressure of the body weight on the skin. PUs are often seen in patients with spinal cord injury and among bed-bound individuals, especially in the elderly population (1). Among hospitalized adult patients worldwide, PU has a prevalence of 12.8% (2), and patients who develop PU have a 3.6 times higher risk of death than those without PU (3). Most PU patients receive conservative treatment consisting of pressure relief and dressing changes, which can last for months to years (4). However, efficient treatment is lacking, primarily due to the limited knowledge regarding the pathophysiology of PU. A minority of PU patients receive reconstructive surgery with resection of the wound and flap coverage of the defect after the failure of conservative therapy. This is an extensive procedure and is unsuitable for elderly individuals and critically ill patients with multiple comorbidities. Additionally, there is a lack of evidence derived from properly conducted randomized controlled clinical trials to support or deny the role of reconstructive surgery in PU treatment (1). Therefore, a deeper understanding of PU pathophysiology and the identification of therapeutic targets are pressing needs.

To our knowledge, we have performed the first full-length single-cell transcriptomic analysis of human PUs in comparison with normal acute wounds (AWs) and skin from matched healthy donors. This dataset can be explored at a free, browsable web portal (https://www.xulandenlab.com/data). The advancement of single-cell RNA sequencing (scRNA-seq) technology has enabled the generation of molecular maps of human tissues at an unprecedented resolution, and this technology has recently been used to characterize human skin (5–8) and related diseases, such as psoriasis (5), atopic dermatitis (9), and melanoma (10), as well as fibroblast heterogeneity (11) and epidermal stem cells (12, 13) in murine wound models. Although rodent wound models exhibit a wound healing procedure similar to that of humans, they cannot fully reflect the complexity of disordered wound healing in humans (14, 15). An in-depth understanding of wound healing mechanisms in humans has the potential to impact wound therapy and diagnosis directly.

Our study revealed significant shifts in epidermal cellular composition and gene expression patterns in PU wound edges compared to AW and uninjured skin. This high-resolution molecular map of human healing and nonhealing wounds will serve as a useful resource for future studies aiming to enhance the understanding of wound healing biology and chronic wound pathogenesis. Among the many previously unknown or poorly characterized cellular and molecular events uncovered by this scRNA-seq analysis, we focused on a novel subset of keratinocytes expressing major histocompatibility complex class II (MHCII), as they were specifically enriched in PUs with worse healing outcomes after reconstructive surgery. We identified IFNγ in PU wound fluid as a major inducer of MHCII expression in keratinocytes. These MHCII^+^ keratinocytes inhibit T cell activation and may be one of the crucial factors in the dysregulated immune response of PU.

## Results

### Characterization of epidermal cell composition of human skin and wounds

To construct a gene expression map of the human wound-edge epidermis at single-cell resolution, we collected samples of chronic nonhealing wounds and nearby intact skin from five PU patients and uninjured skin and day-seven AW from four healthy donors (Figure 1A, Figure S1, Table S1). PU is a highly heterogeneous clinical entity, as it can affect patients of any age and any health condition (1). As a first effort towards the characterization of PU at the single-cell level, we started with a well-defined patient group, i.e., spinal cord injury patients with grade IV PU [according to the European Pressure Ulcer Advisory Panel (EPUAP) classification system] (16) in the gluteal region who were treated with reconstructive surgery. During the operation, we were able to obtain biopsies with sufficient quality and quantity for subsequent molecular analysis without introducing any additional harm to the patients. In addition, we recruited healthy donors who were matched to these PU patients in terms of age, gender, and ethnicity, created surgical wounds at their gluteal area, and then collected wound-edge tissues seven days later, when wound repair was at the proliferative phase (Figure 1A, Figure S1, Table S1) (17).

**Figure 1.**
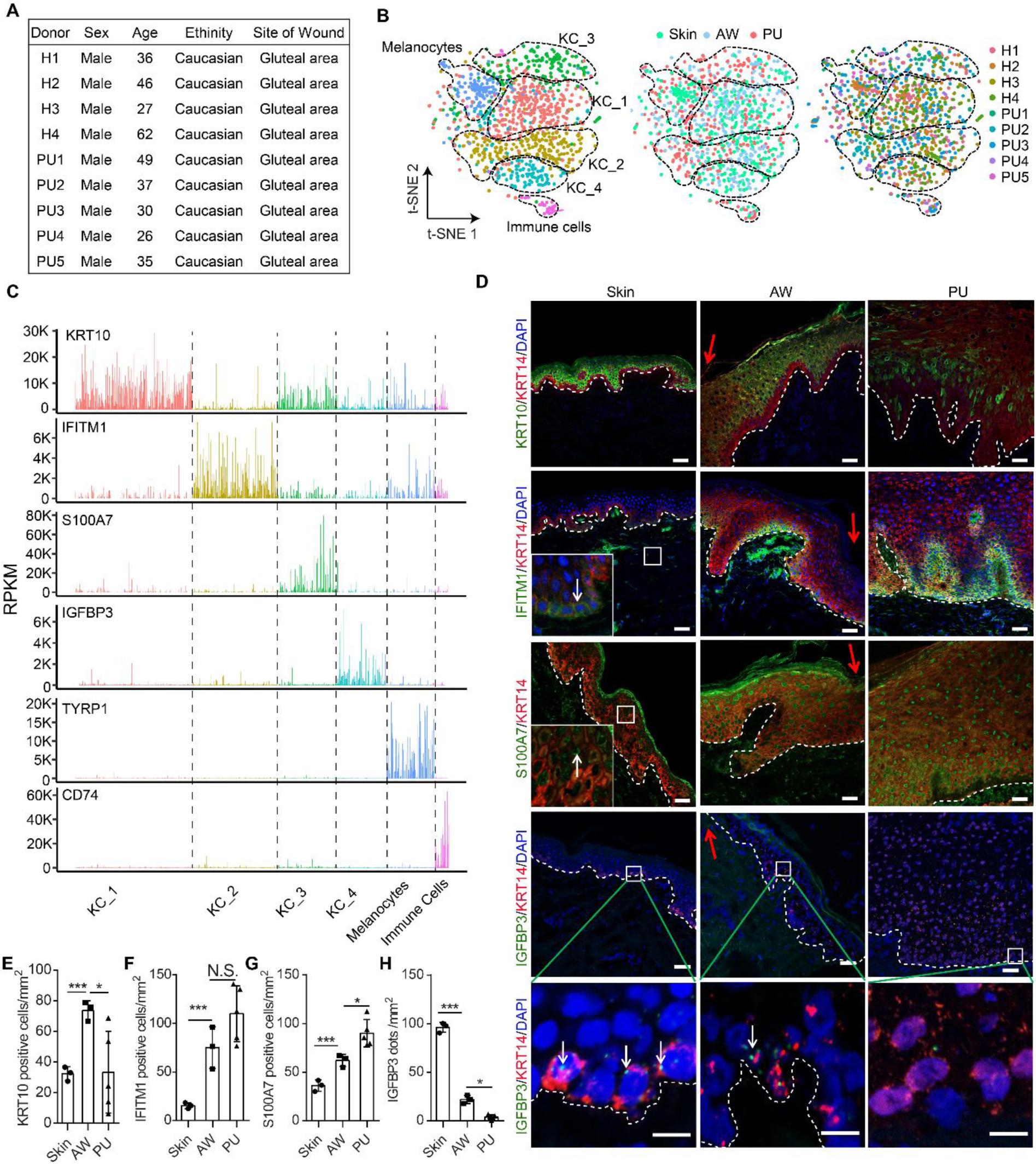
Characterization of epidermal cell composition of human skin and wounds. **A,** Demographics of the subjects analyzed by scRNA-seq. **B**, tSNE projection of all epidermal cells sorted by cell clusters (up), disease conditions (middle), or donors (bottom) (*n* = 1170 cells from 5 PUs, 4 AWs, and 4 intact skin). PU, pressure ulcer; AW: acute wound. **C**, The abundance of selected marker genes of each cell cluster is shown in all the cells. **D,** Immunofluorescence co-staining of KRT14 (red) and KRT10 or IFITM1 or S100A7 (green) in the skin (*n* = 3), AW (*n* = 3), and PU (*n* = 5). In situ hybridization of IGFBP3 (green) and KRT14 (red) mRNAs in the skin (*n* = 3), AW (*n* = 3), and PU (*n* = 5). The red arrows indicate wound edge, while the white arrows highlight a few positively-stained cells. Nuclei were co-stained with DAPI. Scale bar = 50 µm or 10 µm in the enlarged regions. **E**-**H**, KRT10^+^ KRT14^+^, or IFITM1^+^KRT14^+^, or S100A7^+^KRT14^+^, or IGFBP3^+^KRT14^+^ cells were counted and normalized with the area of each field. * p<0.05, *** p<0.001, N.S. not significant; Student’s t-test (**E**-**H**).

We isolated viable epidermal cells and performed Smart-seq2 scRNA-seq, a highly sensitive method for full-length mRNA sequencing at the single-cell level (18). Following stringent quality control, we retained data from 1170 epidermal cells with an average of 5223 genes expressed in each cell (Figure S2A). This number of cells was expected to yield sufficient detection power even if the subtype of interest accounted for only 1% of the epidermal cells (19). The average expression level in scRNA-seq data was highly correlated with the bulk cell RNA sequencing data of the same sample (Spearman correlation coefficient > 0.95), confirming that the single cells we analyzed provided an unbiased representation of all the cell populations in the samples (Figure S2B). These 1170 epidermal cells were segregated into six clusters by Seurat2 clustering (following canonical correlation analysis to minimize batch effects from individual sample variability) (20) (Figure 1B, Figure S2C, D). Each of these clusters contained cells from at least eight out of a total of nine donors and all three sample types, i.e., skin, AW, and PU (Figure 1B). According to the expression of established markers, we annotated one melanocyte cluster (TYRP1^+^, n=149), one immune cell cluster (CD74^high^, n=43), and four keratinocyte clusters (KC_1-4, Figure 1C, Figure S3A, B, Dataset S1). A comparison of our data with a recently published human epidermis scRNA-seq dataset revealed that KC_1 (n=367) and KC_3 (n=189) represented spinous and granular keratinocytes, respectively, while both KC_2 (n=269) and KC_4 (n=153) were basal layer keratinocytes (Figure S3C) (5). Next, we selected the top differentially expressed genes of each cell cluster as marker genes, i.e., KRT10 for KC_1, IFITM1 for KC_2, S100A7 for KC_3, and IGFBP3 for KC_4 (Dataset S1). We confirmed the spatial localization of each keratinocyte cluster in human skin and wounds by detecting these markers at the protein level (by immunofluorescence staining, IF) or RNA level (by fluorescence in situ hybridization, FISH) (Figure 1D-H, Figure S3D).

### Molecular features of epidermal keratinocytes

We further charted keratinocyte differentiation status by aligning the cells based on the expression patterns of five established differentiation markers (21). Most KC_2 and KC_4 cells were assigned to the undifferentiated (KRT5/14^+^KRT1/10^-^CALML5^-^) or early differentiation states (KRT5/14^+^KRT1/10^+^CALML5^-^), whereas KC_1 and KC_3 cells were enriched with keratinocytes at a more differentiated state (KRT5/14^+^KRT1/10^+^CALML5^+^) (Figure 2A-C). Additionally, analysis of cell cycle-related gene expression revealed more proliferative cells in the KC_2/4 clusters than in the KC_1/3 clusters (Figure 2D-G) (22). Gene Ontology (GO) analysis of the marker genes suggested different roles for each keratinocyte cluster, e.g., lipid metabolism and Ras protein signal transduction for spinous keratinocytes (KC_1) and immune response for granular keratinocytes (KC_3). Both basal layer keratinocyte clusters (KC_2 and KC_4) played a role in extracellular matrix organization, while KC_2 keratinocytes were also involved in the response to external stimuli and stem cell development (Figure 2H, Dataset S2). In the interfollicular epidermis of human skin, stem cells reside at the basal layer and exhibit cell heterogeneity (8). Our study revealed two keratinocyte clusters, KC_2 and KC_4, both of which strongly express the known basal keratinocyte signature, that is, KRT14, ITGA6, and ITGB1 (23, 24) (Figure 2I). However, KC_2 differs from KC_4 in its higher expression of immunomodulatory genes such as IFITM1, IFITM3, CCL2, IL1R2, and TIMP1. Meanwhile, KC_2 has a lower level of IGFBP3, which is expressed exclusively in the basal keratinocytes of the suprapapillary epidermis and inhibits cell proliferation (25, 26) (Figure 2I. Dataset S3).

**Figure 2.**
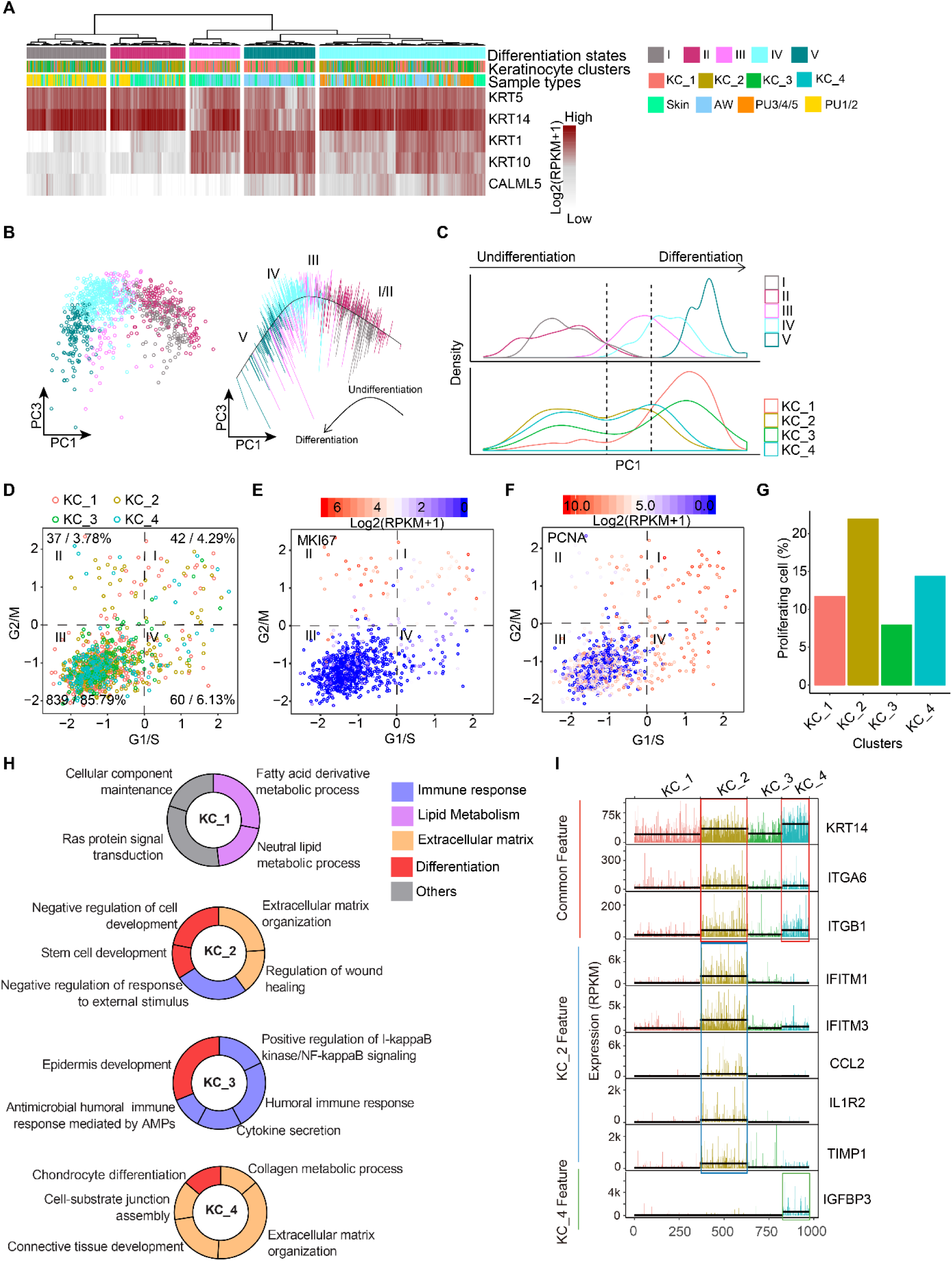
Molecular features of keratinocyte clusters. **A**, Hierarchical cluster analysis of differentiation marker gene expression in keratinocytes analyzed by scRNA-seq. **B,** The expression of these differentiation markers was projected onto PCA, and the principle curve shows the differentiation trajectory. **C**, Cell densities along PC1 were shown for cells in each differentiation stage (upper panel) or cluster (lower panel). **D**, Cell cycle scores of the G1/S and G2/M phases were calculated for each keratinocyte. The expression levels of MKI67 (**E**) and PCNA (**F**) were shown in the cell cycle signature plots. **G,** Proportion of proliferating cells (cells in the quadrants I, II, IV of the cell cycle signature plot) in each keratinocyte cluster. **H**, Gene ontology (GO) analysis was performed for the top 100 upregulated genes of each cluster compared to the rest of the keratinocyte clusters. The donut charts show the percentage of the genes associated with the GO term, and the top GO terms with the lowest p-values are depicted. **I**, Comparison of gene expression between KC_2 and KC_4: the abundance of a few selected genes are shown in all the keratinocytes analyzed by scRNA-seq.

This integrated view of epidermal cells from healthy skin, AW, and PU allowed us to compare the cellular composition and gene expression in homeostatic skin and healing and nonhealing wounds and, importantly, to explore the molecular basis of the pathological changes in PU.

### Epidermal cell heterogeneity in acute wounds and pressure ulcers

Compared to healthy skin, we found fewer melanocytes at the wound edges of AW and PU (Figure 3A, B), in line with previous observations in burn injuries; this phenomenon has been attributed to inflammation-induced cell destruction (27). In contrast, more epidermal immune cells were detected in the PU than in the healthy skin or AW (Figure 3A, B). We compared our results with a published scRNA-seq dataset of 236 epidermal immune cells (5). We identified three hematopoietic subsets in our samples, i.e., CD207^+^CD1A^+^ Langerhans cells (LCs), CD1C^+^CD301A^+^ myeloid dendritic cells (DCs), and CD3^+^ T cells, among which epidermal DCs and T cells were mainly detected in PU (Figure 3C-E, Figure S3E). Comparing AW and skin keratinocytes, we observed increased frequencies of spinous (KC_1) and granular (KC_3) keratinocytes during wound repair. Moreover, we detected fewer spinous (KC_1) and basal layer (KC_4) keratinocytes and more granular (KC_3) keratinocytes in the PU than in healthy skin or AW (Figure 3A, B, Figure 1C-G, Figure 3F).

**Figure 3.**
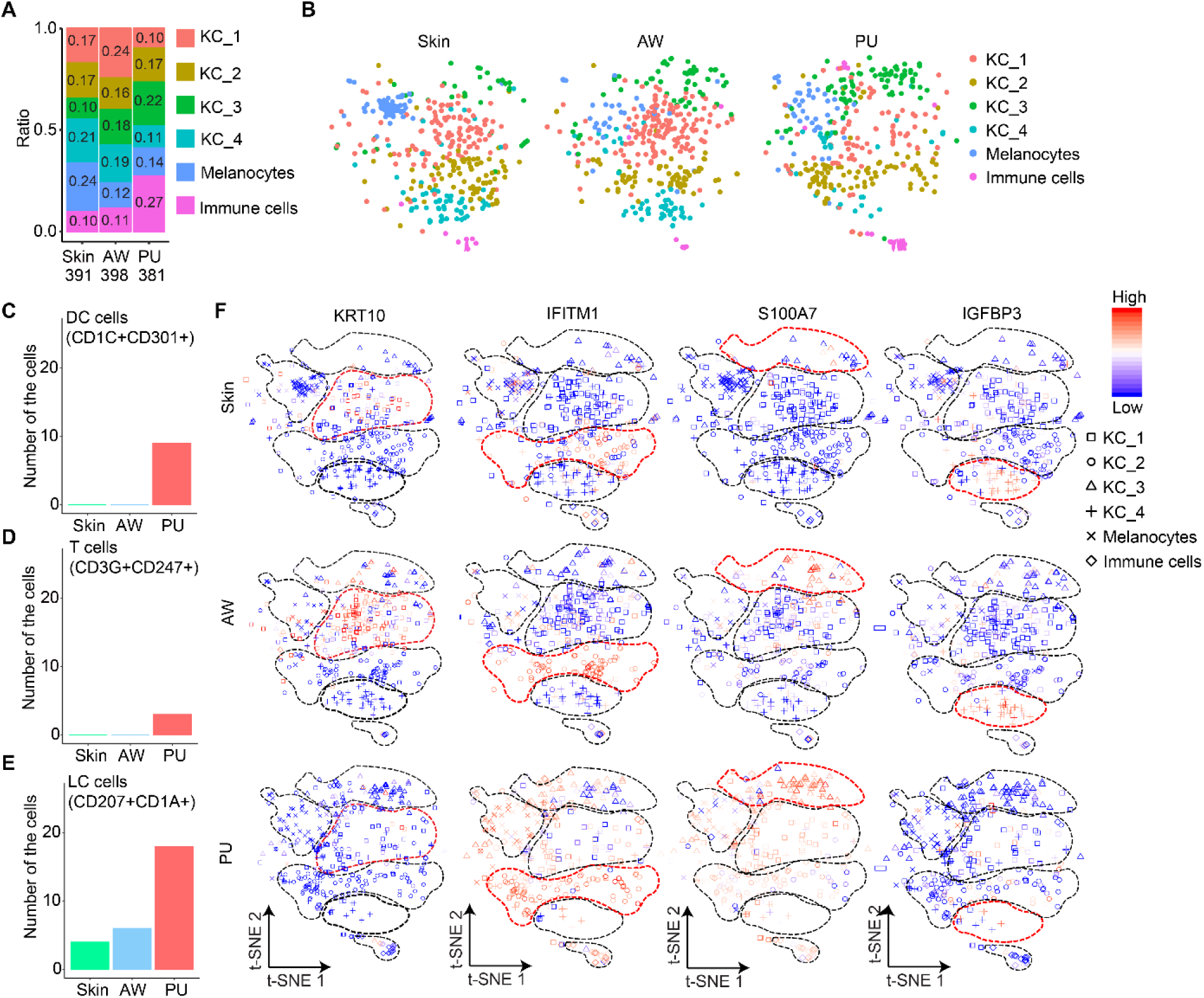
Altered epidermal cell heterogeneity in acute wounds and pressure ulcers. **A**, Frequency distribution and **B**, tSNE projections of the six epidermal cell clusters in the skin (*n* = 391 cells), AW (*n* = 398 cells), and PU (*n* = 381 cells). **C-E**, The number of immune cells identified by scRNA-seq, including CD207^+^CD1A^+^Langerhans cells (LC), CD1C^+^CD301A^+^ myeloid dendritic cells (DC), and CD3^+^ αβ T cells, in the skin, AW, and PU samples. **F,** t-SNE overlay of marker gene expression in the skin, AW, and PU scRNA-seq datasets.

### Differential gene expression in epidermal cells of acute wounds and pressure ulcers

Next, we compared the gene expression in the same cell types from the healthy skin, AW, and PU samples. Unlike melanocytes, which share similar gene expression profiles between AW and PU (Figure 4A, Figure S4A, Dataset S4), keratinocytes exhibited unique, contrasting molecular signatures in PU and AW (Figure 4B, Figure S4B, Dataset S4). GO analysis revealed that genes involved in neutrophil-mediated immunity [e.g., FABP5 (28), S100A7, S100A8, and S100A9 (29)] were strongly upregulated in PU keratinocytes compared to keratinocytes from AW or uninjured skin. However, during normal wound healing (AW vs. skin), the expression of these genes was only slightly enhanced (Figure 4C, Figure S4B, Dataset S5). In addition, the expression levels of genes essential for cellular homeostasis of transition metals, in particular, zinc ions [e.g., MT2A (30), MT1E, FTH1, and FTL (31)], were higher in PU keratinocytes than in AW keratinocytes (Figure 4C, Figure S4B, Dataset S5). The potential role of zinc signaling in chronic wound pathology is supported by recent studies showing that zinc deficiency exacerbates PU by increasing oxidative stress and ATP levels in mouse skin (32), and successful treatment of human venous ulcers changed zinc ion homeostasis in the wound bed (33). Additionally, compared to keratinocytes from AW and uninjured skin, PU keratinocytes expressed higher levels of apoptosis-related genes [e.g., DUSP1 (34) and RHOB (35)], which may be related to the massive activation of the unfolded protein response (e.g., HSPH1 and DNAJB1 (36)) (Figure 4C, Figure S4B, Dataset S5). Interestingly, deregulation of the unfolded protein response has been previously implicated in impaired skin wound healing in diabetes (37). The enhanced apoptosis in PU was confirmed by immunostaining, which showed increased cleaved caspase-3-positive cells in the PU samples compared with the skin and AW samples (Figure 4D).

**Figure 4.**
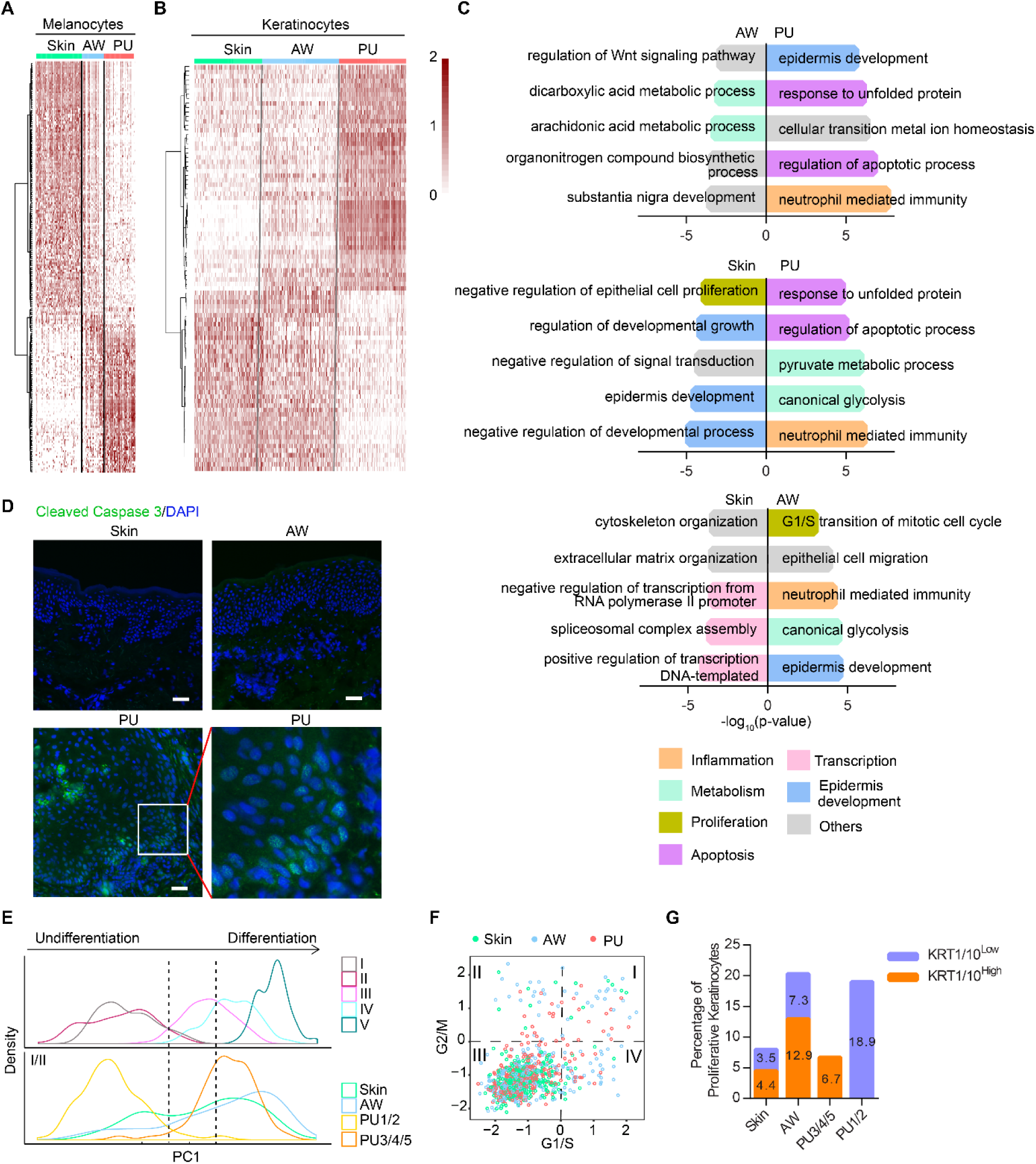
Differential gene expression in epidermal cells of acute wounds and pressure ulcers. Heatmap illustrates the genes differentially expressed in melanocytes (**A**) and keratinocytes (**B**) from the skin, AW, and PU at an individual cell level. **C**, Gene ontology (GO) analysis was performed for the top 150 upregulated genes in comparisons of AW vs. PU (top panel), skin vs. PU (middle panel), and skin vs. AW (bottom panel). The charts show the top five most significant biological process terms in each condition. **D**, Immunofluorescence staining of cleaved Caspase 3 (green) in the skin (n = 2), AW (*n* = 2), and PU (*n* = 5). Nuclei were co-stained with DAPI. Scale bar = 50 µm. **E**, Cell densities along PC1 were shown for cells in each differentiation stage (upper panel) or sample type (lower panel). **F**, Cell cycle scores of the G1/S and G2/M phases were calculated for each keratinocyte in the Skin, AW, and PU. Proliferative cells are the ones in the quadrants I, II, and IV of the cell cycle signature plot. **G**, The percentage of proliferative keratinocytes with low or high KRT1/10 expression was calculated for the skin, AW, and PU samples.

We further compared keratinocyte differentiation and proliferation in the skin, AW, and PU based on the expression of related marker genes. Along the differentiation trajectory reconstructed with pseudotime analyses (Figure 2B), we found that the proportion of differentiated keratinocytes was higher in AW than in the skin (Figure 4E). In contrast to the skin and AW keratinocytes that were distributed along the differentiation axis, PU keratinocytes clustered in differentiation state I (for donors PU1 and PU2) or IV (for donors PU3, PU4, and PU5), suggesting a dysregulation of the differentiation program (Figure 4E). Abnormal keratinocyte differentiation has also been reported in other types of chronic wounds, e.g., venous ulcers and diabetic foot ulcers (38, 39). Furthermore, we identified proliferating keratinocytes based on their expression of cell cycle-related genes (Figure 4F) (22). Interestingly, we found that a significant proportion of these proliferating cells expressed the early differentiation markers KRT1 and KRT10 (Dataset S6), confirming previous findings showing significant proliferative and tissue-regenerative capacity of human early differentiating keratinocytes (40). We detected more proliferating keratinocytes in AW than in skin (Figure 4G). However, in donors PU3, PU4, and PU5, all proliferating keratinocytes were KRT1/10^high^, whereas all proliferating keratinocytes in PU1 and PU2 were KRT1/10^low^ (Figure 4G). Notably, in PU3, PU4, and PU5, but not PU1 and PU2, the proportion of proliferating cells was lower than that in the AWs (Figure 4G). This finding not only revealed dysregulated keratinocyte differentiation and proliferation in PU tissue but also suggested that PU exhibited a higher degree of inter-individual variability than uninjured skin or AW.

### Stratification of PU into two subtypes with distinct molecular features correlating with clinical outcomes

To stratify the PU patients, we performed principal component analysis using the 4000 most variable genes on all PU keratinocytes, which separated PU3, PU4, and PU5 (hereafter named ‘PU group 1’, PU_G1) from PU1 and PU 2 (‘PU group 2’, PU_G2) (Figure 5A). Interestingly, GO analysis of the differentially expressed genes (DEGs) revealed that the PU_G1 keratinocytes expressed genes involved in MHCII-mediated antigen presentation (Figure 5B, Datasets S7, and S8). Following this observation, we discovered a subset of keratinocytes expressing the canonical components of the MHCII machinery (e.g., CD74 and HLA-DRB), which are typically detected in professional antigen-presenting cells (APCs), such as Langerhans cells in the epidermis (Figure 5C). The presence of MHCII on keratinocytes was unlikely to be due to doublets of keratinocytes and APCs, as the MHCII^+^ and MHCII^-^ keratinocytes had similar mean gene expression and ERCC spike-in RNA was used as control (Figure S5A, B). Next, we costained KRT14 and HLA-DR mRNAs using FISH and proteins using IF in additional clinical samples. We confirmed that MHCII^+^ keratinocytes were overrepresented in PU_G1 compared to PU_G2, AWs, or the skin, as shown by scRNA-seq analysis (Figure 5D, E, Figure S5C, K, Supplementary Video 1). On the other hand, the PU_G2 keratinocytes expressed higher levels of pro-proliferation and pro-angiogenesis genes than the PU_G1 keratinocytes (Figure 5B, Dataset S8). We confirmed the GO analysis results by IF costaining of KRT14 and the proliferation marker Ki-67 (Figure 5E). We detected more proliferative keratinocytes in PU_G2 than in PU_G1, which was also in line with previous cell cycle analysis results (Figure 4G). Moreover, scRNA-seq analysis showed that there were more epidermal immune cells in the PU_G1 than in the PU_G2 samples (Figure S5D-G). Additionally, IF staining revealed a higher density of T cells in PU-G1 ulcers than in PU-G2 ulcers, 5-fold in the epidermis and 3-fold in the dermis (Figure S5H-J).

**Figure 5.**
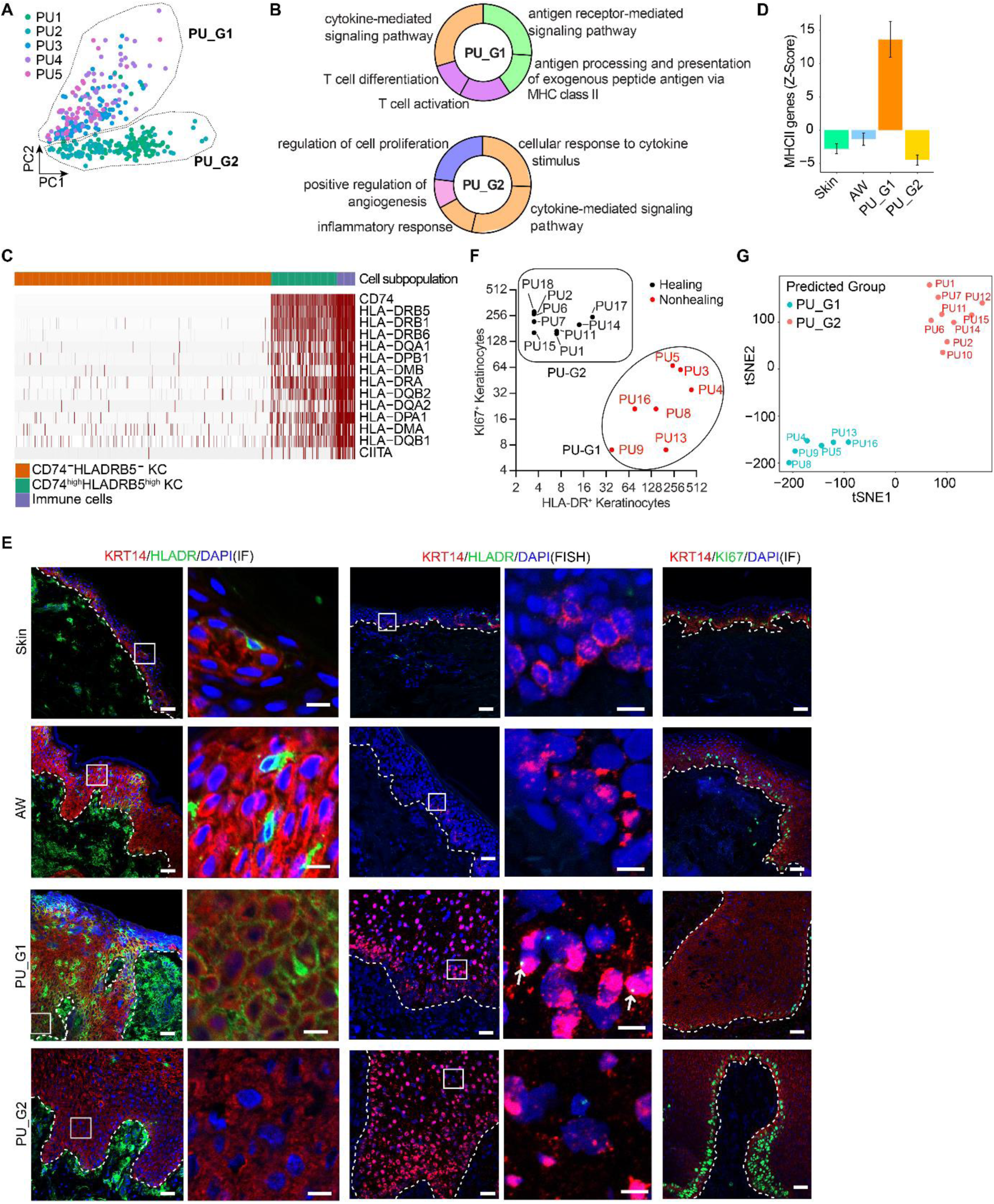
Stratification of PU into subtypes with distinct molecular features correlating with clinical outcomes. **A**, Principle component analysis of keratinocytes from PUs according to their expression of 4000 most variable genes. **B**, Gene ontology (GO) analysis of the top 100 differentially expressed genes between group 1 (G1) and group 2 (G2) PU keratinocytes. The donut charts show the percentage of the genes associated with the GO term, and the top five most significant terms are depicted. **C**, Heatmap shows the expression of MHCII-related genes in CD74^-^HLA-DRB^-^ keratinocytes (KC), CD74^high^HLA-DRB^high^ KC, and immune cells. **D**, The abundance of MHCII-related gene expression is shown in the keratinocytes from the skin, AW, and the two groups of PUs. **E**, Immunofluorescence co-staining (IF) of KRT14 (red) and HLADR (green) or KI67 (green) in skin (*n* = 3), AW (*n* = 3), PU G1 (*n* = 7) and G2 (*n* = 9). In situ hybridization of KRT14 (red) and HLA-DRA (green) mRNAs was performed in tissues from the same donors. Nuclei were co-stained with DAPI. Scale bar = 50 μm or 10 μm in the enlarged regions. **F**, HLADR^+^ and KI67^+^ keratinocytes (identified by KRT14 staining) were counted in 16 PU wound-edge samples analyzed by IF (Figure S6). One month after the surgery, the PUs healed or not healed are indicated with black and red dots, respectively. **G,** tSNE projection of the pressure ulcer samples based on the QRT-PCR analysis of CIITA, CD74, CIITA and VEGFA (Figure S7A-D).

To validate these findings, we performed IF staining of MHCII and Ki-67 in wound-edge biopsies from 16 PU patients treated with reconstructive surgery. We found seven PUs enriched with MHCII^+^ keratinocytes but lacking proliferating keratinocytes (PU_G1), whereas nine PUs had few MHCII^+^ keratinocytes but active proliferating keratinocyte (PU_G2) (Figure 5F, Figure S6). The healing outcomes in these 16 patients were evaluated one month after the operation independently by two surgeons who had not seen the experimental data associated with these patients. Wounds with closed edges and without signs of infection, exudate, or necrotic tissue were diagnosed as healed wounds. Interestingly, all the patients in PU_G2 were identified as ‘healed wounds’; however, none of the patients in PU_G1 fulfilled these criteria of healing (Figure 5F, Table S1). Moreover, based on the qRT-PCR analysis of wound biopsies for the expression of CD74, CIITA, HLA-DRA, and VEGFA, which are among the top DEGs identified in scRNA-seq between PU-G1 and PU-G2, the ‘healing’ and ‘nonhealing’ PUs could also be separated in the tSNE plot (Figure 5G and Figure S7A-D). Notably, we also checked other clinical parameters, including patient age, wound size and duration, circulating C-reactive protein and leukocytes, and bacterial colonization and infection, but did not find that any of them could be used to distinguish the two groups of PUs with varied healing outcomes (Table S1, Figure S7E-I). Despite the small patient cohort, our study provides proof of the principle that PU can be stratified into subtypes with distinct molecular features correlating with clinical outcomes.

### The formation and function of MHCII^+^ keratinocytes in pressure ulcers

To understand the mechanism triggering MHCII expression in keratinocytes, we performed gene set enrichment analysis (GSEA) for the DEGs between the MHCII^+^ and MHCII^-^ keratinocytes, which revealed an upregulation of IFNγ signaling in MHCII^+^ keratinocytes (Figure 6A). In line with this, we found that stimulation of human primary keratinocytes with IFNγ strongly induced the expression of MHCII and its transactivator CIITA at the mRNA level (Figure 6B). Additionally, we observed increased HLA-DR protein expression in IFNγ-treated keratinocytes (Figure 6C). Furthermore, we showed that the cell-free wound fluid from PU_G1 patients, but not PU_G2 patients, significantly induced keratinocyte expression of CD74 and HLA-DRB (Figure 6D, E, and Figure S8A). Importantly, this effect was blocked by neutralizing IFNγ in the wound fluids, suggesting that IFNγ may account for MHCII expression in PU keratinocytes (Figure 6D, E). Of note, we observed that the application of the IFNγ antibody 24 hours after wound fluid treatment could not reverse CD74 and HLA levels in the cultured keratinocytes (Figure S8B-C). Moreover, GSEA of scRNA-seq data revealed higher expression of IFNγ response-related genes in PU_G1 keratinocytes than in PU_G2 keratinocytes (Fig 6F). qRT-PCR analysis showed that IFNγ expression was increased in PU_G1 compared to PU_G2 full-depth wound biopsies. The levels of IFNγ and MHCII were positively correlated in human skin and wounds in vivo (Figure 6G, H, Figure S8D, E). Together, our data suggest that IFNγ is a major inducer of keratinocyte-intrinsic MHCII expression in human PU.

**Figure 6.**
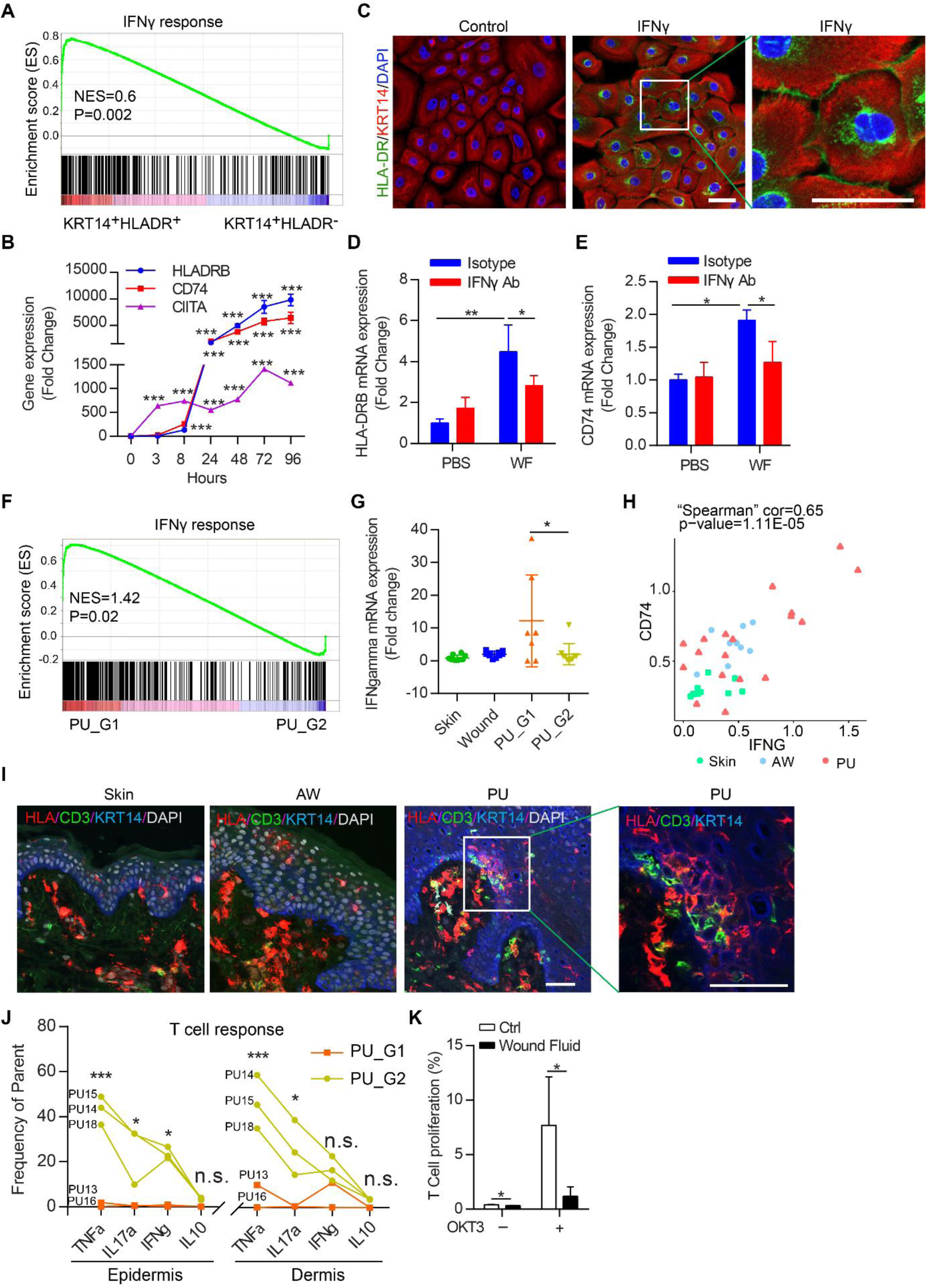
The formation and function of MHCII^+^ keratinocytes in pressure ulcers. **A**, GSEA of the IFNγ-response-related genes among the genes differentially expressed between CD74^high^HLA-DRB^high^ and CD74^-^HLA-DRB^-^ keratinocytes. **B**, qRT-PCR analysis of HLADRB, CD74, and CIITA expression in human adult primary keratinocytes (HEKa) treated with IFNγ (20ng/mL) for 3-96 hours. **C**, Immunofluorescence (IF) co-staining of KRT14 (red) and HLA-DR (green) in HEKa treated with IFNγ (20ng/mL) for 48 hours. **D, E**, HEKa were treated with PU wound fluid (WF, 200μg/mL) supplemented with IFNγ neutralization antibody or isotype control. HLADRB and CD74 expression were detected by qRT-PCR. **F**, GSEA of the IFNγ-response-related genes among the genes differentially expressed between the group1 (G1) and the group2 (G2) PU keratinocytes. **G**, qRT-PCR analysis of IFNG in the full-depth biopsies of human skin (*n* = 10), AW (*n* = 10), group1 (*n* = 7) and group2 PUs (*n* = 10). **H**, Spearman’s correlation analysis between IFNG and CD74 expression detected by qRT-PCR in the above samples. **I**, IF co-staining of HLADR (red), CD3 (green), KRT14 (blue), and DAPI (white) in the skin, AW, and PU. Scale bar = 50µm. **J**, Flow cytometry analysis of TNFα, IL17a, IFNγ, and IL10 production in CD3^+^ T cells from wound-edges of two PU groups. **K**, Keratinocytes were pretreated with wound fluids and co-cultured with autologous T cells activated with OKT3 (*n* = 3 donors). T cell proliferation was analyzed by flow cytometry five days later. The data are presented as mean ± s.d. in (**B**, **D**, **E**, **G**, **K**). * p<0.05, ** p<0.01, *** p<0.001; Student’s t-test.

Next, we explored the potential role of MHCII-expressing keratinocytes in PU pathology. GSEA of scRNA-seq data revealed that MHCII^+^ keratinocytes expressed more antigen presentation- and processing-related genes than MHCII^-^ keratinocytes (Figure S8F). In line with this, we found that MHCII^+^ keratinocytes were close to T cells in PU wound edges by confocal imaging (Figure 6I). Surprisingly, flow cytometry analysis showed that αβ T cells in the PUs with MHCII^+^ keratinocyte overrepresentation (PU_G1) indeed produced fewer inflammatory cytokines, e.g., TNFα, IL-17a, and IFNγ, than the PUs lacking MHCII^+^ keratinocytes (PU_G2, Figure 6J, Figure S9). Moreover, we detected increased epidermal γδ T cells in the PU compared with the skin by flow cytometry (Figure S10A). Similar to αβ T cells (Fig 6J), γδ T cells from PU_G1 also produced less IFNγ than those from PU_G2 (Figure S10B).

To directly study whether MHCII^+^ keratinocytes may impact T cell activation, we cocultured human primary keratinocytes with autologous T cells. We found that pretreatment of keratinocytes with PU-derived wound fluids reduced the concentration of several cytokines important for T cell recruitment and activation, e.g., IFNγ, IL-9, IL-21, GM-CSF, CXCL1, CXCL8, CCL4, and CXCL12, in cell culture supernatants (Figure S10C and D). Moreover, we assessed T cell proliferation by dye dilution. We found that T-cell receptor-induced T cell proliferation decreased from 7.6% to 1.2% (P = 0.03) by coculturing with keratinocytes pretreated with PU1_G1-derived wound fluid (Figure 6K and Figure S10E). These results suggested that keratinocytes might function as atypical APCs and exert an inhibitory effect on T cell activation in PU wound edges.

## Discussion

PU is one of the most frequent causes of death in elderly and wheelchair- or bed-bound individuals (4). Despite the high mortality of patients, the molecular pathology of PU remains mostly elusive, which hampers the development of more effective treatment. For this, we generated the first cellular landscape of human wound-edge tissues at the single-cell level, charting the differences in cell composition and molecular state between nonhealing PU, healing AW, and uninjured skin. Our study focused on the most severe PUs in the gluteal area. It would be interesting to further validate these findings in PU of varying severity and patients with different comorbidities to obtain a complete picture of this complex clinical entity. Nevertheless, this study took an essential first step to gain a single-cell view of the molecular pathology of PU based on a well-defined patient cohort. We made this scRNA-seq dataset of human normal and chronic wounds freely and easily accessible through a web portal (https://www.xulandenlab.com/data) to provide the research community with a unique resource to further explore chronic wound pathology.

The integrated view of cells from healthy skin, AW, and PU allowed us to compare the cellular composition and the gene expression of the same cell type under healthy and disease conditions. Comparing PU with AW, we found that most pathological changes occurred in keratinocytes and immune cells but not in melanocytes, as melanocytes in the healing and nonhealing wounds exhibited similar cellular proportions and transcriptomes. At the PU wound edge, suprabasal KC_3 and basal KC_2 keratinocytes predominated, whereas suprabasal KC_1 and basal KC_4 keratinocytes were lacking. Interestingly, both the KC_3 and KC_2 clusters had gene signatures associated with immune functions. This cellular composition shift was also reflected in the gene expression profile of PU keratinocytes, revealing their intense inflammatory response.

Additionally, a sharp increase in epidermal T cells, myeloid dendritic cells, and Langerhans cells was detected in PU but not in AW, which further advanced our view of the different immune microenvironments between healing and nonhealing wounds. However, due to the low number of immune cells captured in this study, it was challenging to partition them further or characterize their states in more detail. In future studies, pre-enrichment with the CD45 antibody will enable a higher-resolution analysis of immune cells with very low abundance in the epidermis. Moreover, as immune cells reside primarily in the dermis, further study of human wound dermal compartments with single-cell technology will complete the picture regarding the complex immune environments in human chronic wounds.

Here, we discovered a novel MHCII^+^ keratinocyte population that is overrepresented in PUs with worse healing outcomes. Keratinocytes are known to play an active role in skin immunity and inflammation by producing many cytokines and chemokines (17). Our findings suggest that they may also directly interact with T cells via antigen presentation. Our in vitro study showed that such interactions reduced T cell activation; this conclusion was further supported by ex vivo data revealing that T cells from PUs enriched with MHCII^+^ keratinocytes produced fewer inflammatory cytokines. Therefore, we postulated that MHCII^+^ keratinocytes might interfere with T cell function in PU, which was endorsed by previous studies showing that keratinocyte-T cell interactions could lead to T cell anergy or tolerance (41, 42). Additionally, epidermal T cells isolated from human chronic wounds have been shown to be less responsive to stimulation than T cells from acute wounds (43). Together, our findings support the recently raised hypothesis that chronic wound inflammation is persistent but ineffective in combatting infection and healing wounds (43, 44). Interestingly, a small fraction of keratinocytes expressing MHCII was also found in humans (45) and the mouse epidermis under homeostasis (46). In mouse skin, MHCII^+^ keratinocytes were shown to control homeostatic type one responses to the microbiota (46). Thus, MHCII expression by keratinocytes occurs in both physiological and pathological conditions but may have different consequences in these different contexts.

Our study suggests the potential significance of IFNγ in human PU pathology by identifying it as a major inducer of MHCII^+^ keratinocytes and associating its upregulation in PUs with delayed healing. IFNγ has been shown to induce MHC class II expression in keratinocytes (46–50) and to inhibit keratinocyte proliferation (51). In rodent models, IFNγ has also been found to inhibit angiogenesis and collagen deposition, thus hampering wound repair (52, 53). Based on these pieces of evidence, it would be tempting to test whether the local blockage of IFNγ signaling may improve wound healing in PU patients. Under this paradigm, humanized anti-IFNγ antibody, which is under development to treat Crohn’s disease, might be repurposed for wound therapy (54).

The high patient-to-patient variability in PU underscores the need for personalized wound treatments. However, biomarkers to stratify subsets of nonhealing patients and to guide therapy are lacking (55). Our study showed that even within a group of patients with similar clinical manifestations, the gene expression profiles of PU wound-edge keratinocytes could distinguish between different subtypes. More interestingly, those patients with MHCII^+^ keratinocyte overrepresentation accompanied by Ki-67^+^ keratinocyte reduction at their wound edges had worse healing outcomes after operation. Despite the limited number of patients included, our study provides a proof of principle that PU can be stratified into subtypes with distinct molecular hallmarks highly correlated with clinical outcomes, underscoring the importance of molecular wound diagnosis. Further validation of our findings in larger patient cohorts may lead to the identification of biomarkers for early recognition of PU patients who may require attentive therapeutic interference. Thus, our study provides insights for future precision medicine in wound care.

## Methods

### Human wound samples

We enrolled 25 healthy donors and 18 PU patients at the Karolinska University Hospital (Stockholm, Sweden) (Table S1). All the clinical materials were taken after the patients’ consent, and the study was approved by the Stockholm Regional Ethics Committee and conducted according to the Declaration of Helsinki’s principles.

For the healthy donors participating in the study of the human in vivo wound model (H1-H14, Table S1), the exclusion criteria were skin diseases, infections, diabetes, unstable heart disease, immune suppression, and bleeding disorder. Using a 4 mm biopsy punch, the surgeon made two full-thickness excisional wounds on the skin of the gluteal area of each donor, and the excised skin was saved as intact skin control. At a re-visit seven days later, the wound-edges were collected using a 6 mm biopsy punch (Figure S1A). The local carbocain-adrenalin injection was used for anesthesia while sampling.

Also, 20mL venous blood and discarded skin were collected from three donors who underwent abdominoplasty (H15-H25, Table S1).

Wound-edge samples and nearby intact skin were collected from patients with grade IV PU (according to the EPUAP classification system) during reconstructive surgery (16). One month after the operation, wound healing outcomes were evaluated at the rehabilitation clinic: wounds with closed edges and without signs of infection, exudate, or necrotic tissue were diagnosed as healed wounds. We confirmed the histology of skin and wound biopsies by hematoxylin and eosin staining (Figure S1C).

During dressing change, PU wound fluids were collected from the gauzes in direct contact with wound beds for two to three days. The gauzes were soaked in 10 mL phosphate-buffered saline (PBS) and centrifuged for 5 minutes at 10000 rpm. The supernatants were passed through filters with 0.22 μm pore size (Sarstedt). Their protein concentrations were then determined using Pierce™ BCA Protein Assay Kit (ThermoFisher Scientific) according to the manufacturer’s instructions.

### Single-cell isolation and sequencing

Skin and wound samples were incubated in Dulbecco’s Modified Eagle Medium (DMEM) containing 5U/ml Dispase II and 1% Penicillin-Streptomycin (ThermoFisher Scientific) at 4 ℃ overnight. The epidermis was separated from the dermis and then digested with 0.025% Trypsin/ EDTA (ThermoFisher Scientific) for 10 minutes at 37 ℃. The cells were washed with DMEM once and then resuspend in 1mL DMEM containing 5μl Calcein (ThermoFisher Scientific) to check cell viability. 1511 viable cells were picked manually by mouth pipetting and then transferred into Smart-seq2 lysis buffer. After single-cell picking, the rest of the cells of each sample were collected in a pellet and lysed in Trizol to extract total RNA. After confirming the good quality of total RNA by Nanodrop, 1ng of total RNA per sample was used for library preparation. cDNA libraries of single-cell and bulk cells were generated using the Smart-seq2 method(18) (Figure S1A). SuperScript II Reverse Transcriptase (Thermo Fisher Scientific) was used in reversed transcription of polyA (+) RNAs, and the strand-switch reaction was performed in second-strand synthesis. KAPA HiFi HotStart PCR Kit (Kapa Biosystems) was used to amplify the cDNA. The cDNA libraries were purified by carboxylated magnetic beads, then their quality and quantity were confirmed by using the Agilent 2100 Bioanalyzer. The sequencing libraries were generated using Tn5 transposase (18) and Nextera XT dual index primers (Illumina) and then performed on the HiSeq 2500 system.

### Single-cell RNA-sequencing data analysis workflow

#### Quality control of scRNA-seq data

All reads were mapped to homo sapiens genome GRCh38 by using the STAR method with default parameters, and uniquely mapped reads were used in further analysis(56). Reads per kilobase million (RPKM) were calculated by using “rpkmforgenes” (57). We applied stringent quality control filters on the sequencing data of each cell, including (1) expression of more than 1000 genes with RPKM > 1; (2) total read counts > 50K; (3) uniquely mapping ratio > 0.4; (4) Spearman correlation coefficients between any two nearest cells > 0.4. 1170 out of 1511 cells passed these four criteria.

#### Unsupervised identification of epidermal cell clusters

The canonical correlation analysis (CCA) was performed with an R package “Seurat” (Version 2.3.4 with default parameters) for the 1170 cells that passed the quality control. This analysis reduced inter-sample or donor variability(20). The cells were projected to the first two t-distributed Stochastic Neighbor Embedding (t-SNE) dimensions by the first 20 CCA dimensions (function “RuntSNE”)(58). We identified six cell clusters at a resolution of 1.0 (function “FindClusters”), and CCA clustering of the same dataset with a lower (0.8) or a higher (1.2) clustering resolution was shown in Figure S2C, D. We selected the resolution of 1.0 based on prior knowledge and comparison with a published scRNA-seq dataset of human skin epidermis (5). Cell types were identified according to canonical and novel markers revealed by differential expression analysis. We calculated the differentially expressed genes (DEGs) for individual cell clusters compared to the rest cell clusters using the method of Likelihood-ratio test for single-cell gene expression (function “FindMarkers” with test.use = bimod, min.pct = 0). The DEGs were expressed in more than 50% cells within the respective cluster and with a log2 (fold-change) >= 0.5 and adjusted p-value < 0.05.

#### Gene ontology (GO) analysis

GO analysis was performed for the DEGs with Enrichr (59, 60). Among the GO biological process terms with p<=0.01 and containing at least four genes from the input gene list in each GO term, the top five (if there were more than five) GO terms with the lowest p-values were shown. GO terms with identical gene lists were combined.

#### Comparison of our dataset with a published scRNA-seq dataset of human skin epidermis

To evaluate the similarities of the cell types identified here with a previous study by Cheng et al. (5), we retrieved the filtered read count matrix of 25129 cells of healthy human trunk epidermis (EGAS00001002927) from the European Genome-phenome Archive. The variable genes that were in common in both our and their datasets were calculated by using the R function “variableGenes” (R package “MetaNeighbor” version 0.100.0). In Cheng et al. ’s work, 11 cell types were identified in human normal and psoriasis epidermis, i.e., immune cells, two melanocyte populations (mel1 and mel2), and eight keratinocyte populations (granular, spinous, mitotic, channel, follicular, WNTI, basal1, and basal 2) (5). The similarities between these 11 cell clusters and the six cell clusters identified in our study were evaluated by the mean area under the receiver operator characteristic curve (AUROC) matrix (R package “MetaNeighbor” version 0.100.0) (61) (Figure S3C). Moreover, we compared the 43 immune cells captured in our study with the 236 immune cells characterized in Cheng et al. ’s study, which allowed us to identify the most similar (AUROC score >=0.8) cell types in our immune cell cluster (Figure S3E).

#### Pseudotime reconstruction of skin and wound keratinocytes based on the expression of differentiation-related genes

Based on the expression patterns of five established differentiation markers, i.e., KRT5, KRT14, KRT1, KRT10, and CALML5 (21), all the keratinocytes were hierarchically clustered into five states spanning from undifferentiation (states I-II: KRT5/14^high^KRT1/10^low^), early (state III: KRT5/14^high^KRT1/10^high^CALML5^-^) to more differentiation (states IV-V: KRT5/14^high^KRT1/10^high^CALML5^+^) (Figure 2A). The cells were projected to PC1 and PC3 in principle component analysis (PCA), and the pseudo order of the cells was the principle curve from the dimensions of PC1 and PC3 (R package “princurve” version 2.1.3). We specified the state I-II as a starting state, while the state V as a terminal state (Figure 2B). Based on the ordering of the cells in PC1, the composition densities of the cells with various differentiation states (states I-V) or from different samples (skin, AW, and PU) or clusters (KC_1-4) were aligned along the main pseudo axis (Figure 2C, Figure 4E).

#### Cell-cycle signature in keratinocytes

To evaluate keratinocytes’ proliferation status, we calculated the cell cycle scores of the G1/S and G2/M phases for each cell, as previously described (62). This score was defined as the average log2-transformed relative expression of the highly correlated cell-cycle genes (Spearman correlation coefficient > 0.4). The relative expression value of each cell cycle gene was calculated as the RPKM of this gene in individual cells minus its average RPKM in all the cells. The list of genes related to the G1/S and G2/M phases were obtained from a previous study (22) (Dataset S9).

#### Differential expression analysis of cells from the skin, AW, and PU

We compared PUs with either AW or the skin for their gene expression patterns in keratinocytes and melanocytes. Using DESeq2 (version 1.22.2 with default parameters), we identified the DEGs with an adjusted p-value < 0.05, fold change ≥ 2 and expressed in more than 50% cells within each group. The group 1 and group 2 PUs were compared with the skin or AW separately, and the DEGs showing up in both comparisons were considered the common DEGs in PUs.

#### Principle component analysis of PU samples

The 381 keratinocytes from the five PU samples were clustered by PCA (R function “prcomp” with default parameters) with the 4000 most variable genes, which were determined with Brennecke’s method (63). The differential expression analysis was performed for the keratinocytes from the group 1 and group 2 PUs using the R package “DESeq2” (version 1.22.2) as described above.

### Statistics

All data were expressed as mean ±s.d. or mean ±s.e.m. and plotted using GraphPad Prism v6. Statistical significance was determined by a two-tailed Student’s t-test or Mann-Whitney U Test. The correlation between the expressions of different genes in the same sample set was made using Spearman’s correlation test on log-transformed data. For all statistical tests, P values < 0.05 were considered to be statistically significant. P values and analysis methods are also described in each figure legend.

### Data and software availability

The raw data and the processed matrices of raw count and RPKM of scRNA-seq analysis have been deposited in the National Center for Biotechnology Information Gene Expression Omnibus (GEO) database (accession no. GSE137897 with a token ‘clyhsecmjxulzkl’). The codes are available at https://github.com/shanglicheng/HumanKeratinocyte. A web resource for browsing the scRNA-seq data can be accessed at our resource webpage https://www.xulandenlab.com/data.

The other detailed materials and methods used for immunofluorescence staining; fluorescence in situ hybridization (FISH); image analysis; cell culture and treatments; RNA extraction and qRT-PCR; analysis of cytokine production of T cells from pressure ulcers; flow Cytometry and autologous keratinocyte-T cell co-culture are provided in SI Appendix, Supplemental Experimental Procedures. Patients’ information is listed in SI Appendix, Table S1. Antibodies, primers, and chemicals used in this study are listed in SI Appendix, Table S2.

## ACKNOWLEDGMENTS

We express our gratitude to all the patients and healthy donors who took part in this study. We thank Dr. Maria Kasper (Karolinska Institutet) and Dr. Stanley Sing Hoi Cheuk (Göteborgs University) for discussion and advice. We thank Dr. Zhuang Liu, Borislav Ignatov (Karolinska Institutet), Hua Zhang, and Dr. Yonglong Dang (Uppsala University) for technical support. We thank Madeleine Stenius (Rehab Station Stockholm Academy) for clinical sample collection. This work was supported by Swedish Research Council (Vetenskapsradet, 2016-02051 and 2018-02557), Ragnar Söderbergs Foundation (M31/15), Hedlunds Foundation, Welander and Finsens Foundation (Hudfonden), Åke Wibergs Foundation, Jeanssons Foundation, Swedish Psoriasis Foundation, Ming Wai Lau Centre for Reparative Medicine, Tore Nilson’s Foundation, Lars Hiertas Foundation, and Karolinska Institutet.

## AUTHOR CONTRIBUTIONS

N.X.L. and Q.D. conceived the study; D.L., Y.P., J.K., E.K.H., M.T., and L.Z. performed the experiments; S.C., D.L., and Y.T.C. performed bioinformatics analysis; X.C. and L.E. assisted with experimental design and data interpretation; P.S. and K.P. collected clinical samples and information; N.X.L. wrote the manuscript, which was commented on by all the authors.

