## Supplemental Figures for "Single-cell analysis reveals MHCII expressing keratinocytes in pressure ulcers with worse healing outcomes"

### Supplemental material

**Single-cell transcriptomics of human pressure ulcers reveals MHCII expressing keratinocytes in patients with a worse healing outcome**

Dongqing Li, Shangli Cheng, Yu Pei, Pehr Sommar, Jaanika Kärner, Eva K. Herter, Maria A. Toma, Letian Zhang, Kim Pham, Yuen Ting Cheung, Xingqi Chen, Liv Eidsmo, Qiaolin Deng, Ning Xu Landén

#### Supplemental experimental procedures

##### Immunofluorescence staining (IF).

Frozen human skin and wound sections (in 10 μm thickness) were fixed in ice-cold 100% acetone for 5 minutes. After blocked with 2.5% horse serum for 30 minutes, the sections were incubated with primary antibodies or isotype control at 4℃ overnight (Table S2). On the second day, the sections were washed three times with PBS and then incubated with secondary antibodies (Table S2) in 2.5% horse serum in the dark at room temperature for 40 minutes. After washed with PBS three times, the sections were mounted with the ProLong™ Diamond Antifade Mountant with DAPI (ThermoFisher Scientific).

##### Fluorescence in situ hybridization (FISH).

The probes for detecting human IGFBP3, HLA-DR, and KRT14 mRNAs were purchased from Advanced Cell Diagnostics (Table S2). FISH was performed on human skin and wound sections (8μm in thickness) by following the manufacturer's instructions (RNAscope® Multiplex Fluorescent Reagent Kit v2, Document Number 323100-USM). Briefly, fresh frozen tissue sections were fixed in cold 4% formaldehyde for 15 minutes. RNAscope® Hydrogen Peroxide was added to the sections and incubated for 10 minutes at room temperature in the HybEZ™ Humidity Control Tray. The sections were then incubated with RNAscope® Protease IV at room temperature for 30 minutes. Next, the sections were incubated with probes (Table S2) for two hours at 40 ℃ in HybEZ™ II Hybridization System. The hybridization signals were amplified via sequential hybridization of amplifiers and labeled probes (AMP1, AMP2, and AMP3). TSA® plus fluorescein or Cyanine 3 was assigned to the C1, C2, or C3 channels. At last, the sections were mounted with ProLong™ Diamond Antifade Mountant with DAPI (ThermoFisher Scientific).

**Image analysis.**

The sections of IF staining and FISH were visualized with LSM800 confocal laser scanning microscope (Carl Zeiss). The positive cells in each section were counted and normalized with the area of the field. Three sections were evaluated for each sample. The results of the skin and AW from three healthy donors and PU from 5-15 donors were shown. Three-dimensional videos were obtained from z-stacks after processing with Imaris image analysis software.

**Cell culture and treatments.**

Human adult epidermal keratinocytes were purchased from ThermoFisher Scientific or isolated from human skin (see below), and cultured in EpiLife medium supplemented with 1X Human Keratinocyte Growth Supplement (HKGS), 100 U/mL of Penicillin-Streptomycin and 60 µM calcium chloride at 37°C in 5% CO2. Third-passage of keratinocytes at 70% confluence were treated with IFNγ (20ng/mL, R&D) for 3, 8, 24, 48, 72, 96 hours. Alternatively, they were treated with PU wound fluids (protein concentration 200 μg/mL) containing IFNγ neutralization antibody or isotype control (200 ng/mL, Invivogen, Table S2) for 24 hours. Keratinocytes were also treated with PU wound fluids (protein concentration 200 μg/mL) for 24 hours and further treat with IFNγ neutralization antibody or isotype control (200 ng/mL, Invivogen, Table S2) for 24 hours.

**RNA extraction and qRT-PCR.**

Human skin and wound biopsies were homogenized in liquid nitrogen using a Mikro-Dismembrator S (Braun Biotech). Total RNA was extracted using the miRNeasy Mini kit (Qiagen) from homogenized skin tissues or cells by following the manufacturer's instructions. Total RNA was reverse transcribed using the Revert Aid™ First Strand cDNA Synthesis Kit

(ThermoFisher Scientific). Expression of specific genes was quantified by PrimeTime qPCR Assay (IDT) on QuantStudio 7 Flex Real-Time PCR System (Invitrogen) and normalized to the expression of 18S ribosome RNA. The quantification of gene expression was determined by the comparative 2ΔΔCT method. The sequences of all the primers used in this study can be found in Table S2.

**Analysis of cytokine production of T cells from pressure ulcers and healthy skin.**

Biopsies from healthy skin and wound-edge tissues were placed into dispase (5U/mL, Thermo Fisher) at 4 °C overnight. The epidermis was separated from the dermis and put into a collagenase III (570U/mL, Worthington) solution with 5ug/mL DNAse (Roche) for 1.5 hours at 37 °C. After passing through 100μm and then 70 μm filters, the epidermal and dermal suspensions were placed in the RPMI medium containing 10% FBS and 1% Penicillin/Streptomycin on 24-well plate overnight. On the next day, half of the samples were stimulated for 5 hours with PMA (50ng/ml, Sigma-Aldrich), Ionomycin (1ng/ml, Sigma-Aldrich) in the presence of brefeldin A (Golgi plug, BD Bioscience) to assess the cytokine production.

**Flow Cytometry.**

Cells were stained with the Fixation/Permeabilization Solution Kit (BD Bioscience). Briefly, the cells were incubated for 20 minutes at room temperature (RT) with surface staining mix in FACS buffer (PBS with 0,5% FBS and 2mM EDTA) containing viability dye (LIVE/DEAD™ Fixable Yellow Dead Cell Stain Kit, Thermo Scientific), after that, they were washed and then fixed for 20 minutes at RT using the BD Cytofix/Cytoperm buffer (BD Bioscience). After washing twice with BD Perm/Wash (BD Bioscience), the cells were incubated at RT for 30 minutes with the intracellular staining mix in BD Perm/Wash buffer, followed by washing and taking up in FACS buffer. Data were acquired by using the LSR Fortessa flow cytometer (BD Bioscience) and analyzed by using FlowJo V10.6.1.

**Autologous keratinocyte-T cell co-culture.**

*Isolation of keratinocytes.*

Ten 6mm skin punch biopsies were incubated in PBS containing 5U/mL Dispase II and 1% Penicillin-Streptomycin (ThermoFisher Scientific) at 4 ℃ overnight. The epidermis was separated from the biopsies, cut into small pieces, and then incubated in 0.025% Trypsin-EDTA (ThermoFisher Scientific) for 15 minutes at 37 ℃. After mixed with the Defined Trypsin Inhibitor (ThermoFisher Scientific), the epidermal cells were filtered through a 70 μm cell strainer. Epidermal cells were cultured in EpiLife medium supplemented with 1X HKGS, 100 U/mL of Penicillin-Streptomycin, 2.50 µg/mL of Amphotericin B (Fungizone, Gibco) and 60 µM calcium chloride (ThermoFisher Scientific) at 37°C in 5% $CO_2$ until 70%-80% confluency.

*Isolation of T cells.*

Peripheral blood mononuclear cells (PBMCs) from healthy donors were prepared using Ficoll (GE Healthcare) density separation and frozen immediately in fetal bovine serum (FBS) containing 10% dimethyl sulfoxide (DMSO) and stored at –80 ℃. The cells were thawed and kept in RPMI medium with 10% FBS and 1% Penicillin/Streptomycin for 24 hours. Then $CD3^+$T cells were isolated using the human Pan T Cell Isolation Kit (Miltenyi Biotec) and labeled with CellTrace™ Violet Cell Proliferation Kit (Thermo Fisher) according to the manufacturers' protocol and washed twice with EpiLife medium.

*Autologous keratinocyte-T cell co-culture.*

Twenty-four hours before co-culture, keratinocytes were treated with 200 μg/mL of wound fluid from pressure ulcer patients. Then 4*10^4 keratinocytes were co-cultured with 2*10^5 T cells in 1mL fresh EpiLife medium without HKGS in 24-well plate. T cells were stimulated without/with soluble ultra-LEAF™ Purified anti-human CD3 Antibody (1ng/mL) (Biolegend) on day1 and day3 of co-culture. On day5, T cells were collected for flow cytometry analysis.

*Multiplex Immunoassay*

Cell culture supernatants (50 μL) collected on day three and day five during autologous keratinocyte-T cell co-culture were analyzed by using Cytokine & Chemokine Convenience 34-Plex Human ProcartaPlex™ Panel 1A (ThermoFisher Scientific) and the Bio-Plex® 200 Systems (Bio-Rad) according to the manufacturers' protocol.

A

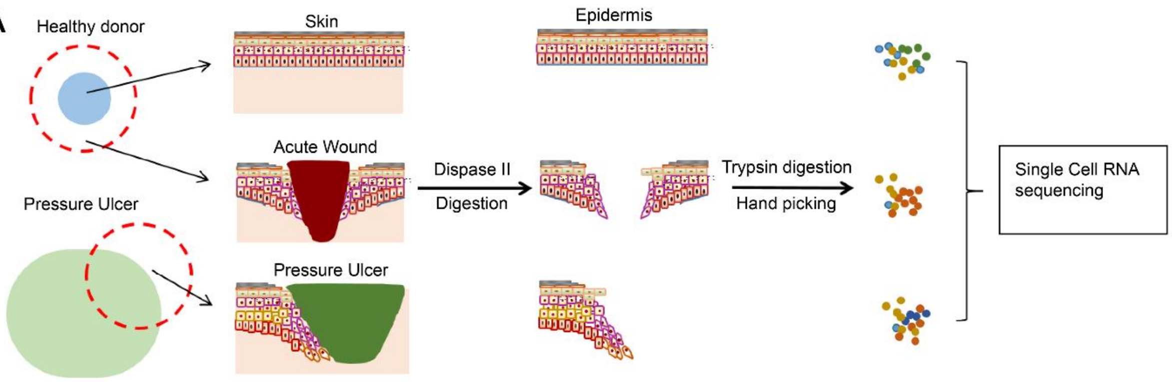


B

| Donor | Age | Gender | Duration of wound (Month) | Wound size (cm) | Diagnosis | Site of wound | Cell number passed QC | | |
|---|---|---|---|---|---|---|---|---|---|
| | | | | | | | Skin | AW | PU |
| H1 | 37 | Male | --- | --- | --- | Gluteal area | 100 | 114 | --- |
| H2 | 30 | Male | -- | --- | --- | Gluteal area | 102 | 82 | --- |
| H3 | 26 | Male | -- | --- | --- | Gluteal area | --- | 117 | --- |
| H4 | 25 | Male | -- | --- | --- | Gluteal area | 107 | 85 | --- |
| PU1 | 36 | Male | 24 | 6x4 | Pressure ulcer | Trochanter majus | --- | --- | 89 |
| PU2 | 46 | Male | 24 | 4x3+4x3 | Pressure ulcer | Groin & Trochanter majus into hip joint | --- | --- | 107 |
| PU3 | 27 | Male | 84 | 2x3 | Pressure ulcer | Tuber ischiadicum | 82 | --- | 81 |
| PU4 | 62 | Male | 24 | 7x3 | Pressure ulcer | Tuber ischiadicum | --- | --- | 61 |
| PU5 | 49 | Male | 18 | 8x5 | Pressure ulcer | Tuber ischiadicum | --- | --- | 43 |

C

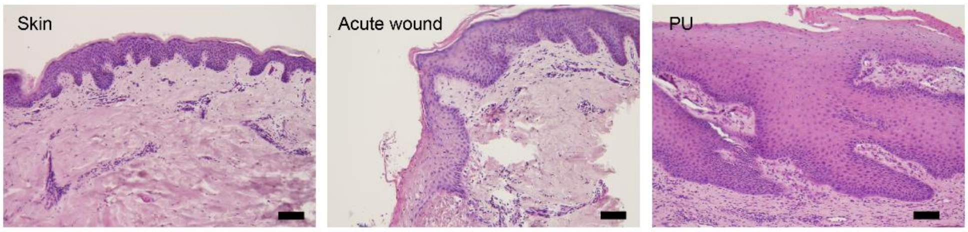


**Figure S1. Single-cell RNA sequencing analysis of human skin and wound-edge epidermis. A,** Schematic overview of the workflow in this study. **B,** Donor information for the biopsies analyzed by scRNA-seq. **C**, Hematoxylin and eosin staining of human skin, acute wound, and pressure ulcer (PU) biopsies. Scale bar, 500 μm.

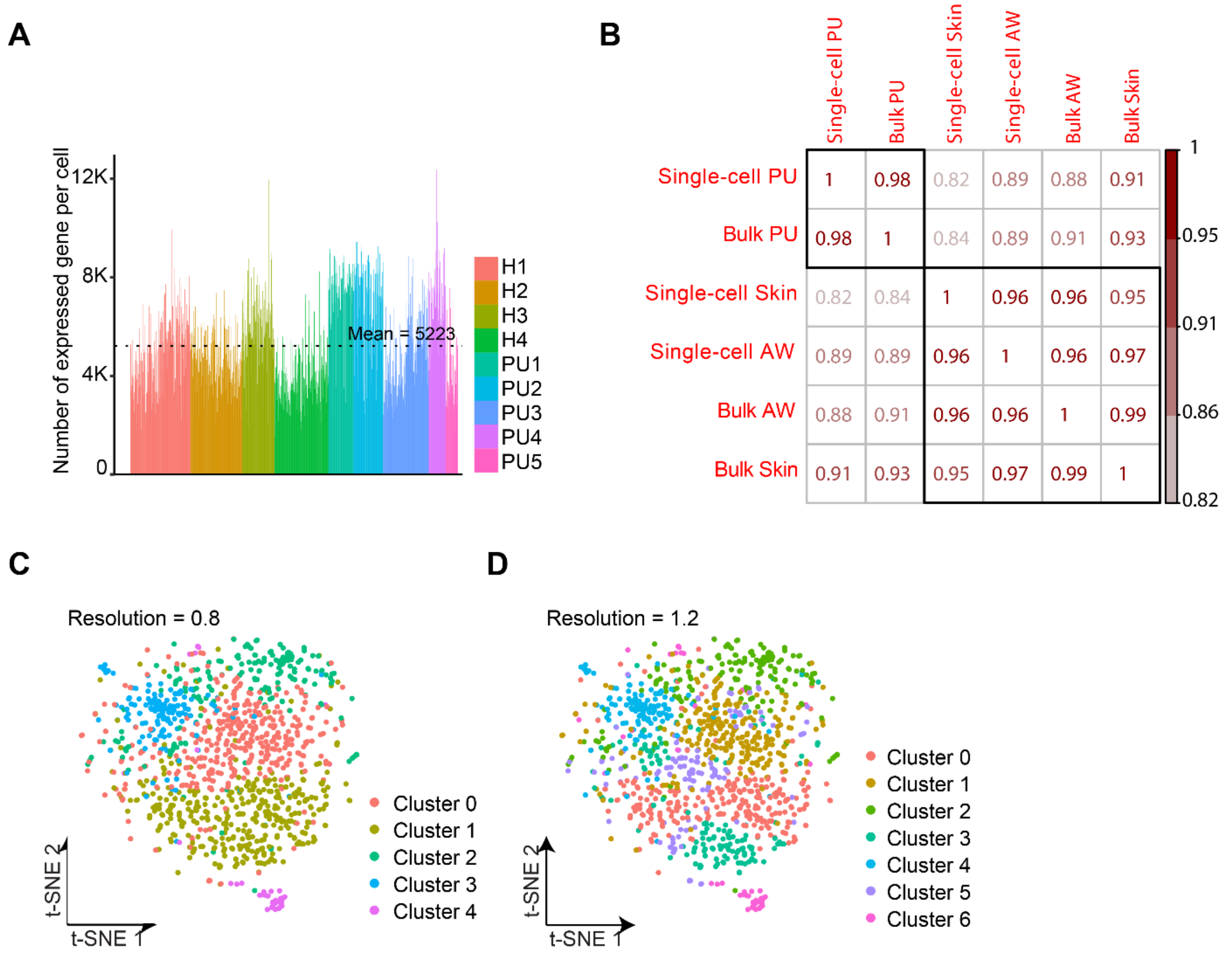


|  | Single-cell PU | Bulk PU | Single-cell Skin | Single-cell AW | Bulk AW | Bulk Skin |
|---|---|---|---|---|---|---|
| Single-cell PU | 1 | 0.98 | 0.82 | 0.89 | 0.88 | 0.91 |
| Bulk PU | 0.98 | 1 | 0.84 | 0.89 | 0.91 | 0.93 |
| Single-cell Skin | 0.82 | 0.84 | 1 | 0.96 | 0.96 | 0.95 |
| Single-cell AW | 0.89 | 0.89 | 0.96 | 1 | 0.96 | 0.97 |
| Bulk AW | 0.88 | 0.91 | 0.96 | 0.96 | 1 | 0.99 |
| Bulk Skin | 0.91 | 0.93 | 0.95 | 0.97 | 0.99 | 1 |

**Figure S2. Canonical correlation analysis (CCA) and Seurat2 clustering of scRNA-seq data. A**, The number of expressed genes (RPKM ≥ 1) in each cell. **B**, Pearson correlation between the average expressions in scRNA-seq data and bulk RNA-sequencing data of human skin, acute wounds (AW), and pressure ulcers (PU). Seurat CCA clustering of all the cells from the skin, AW and PU with a lower (**C**) or higher (**D**) resolution than the clustering resolution in Figure 1B.

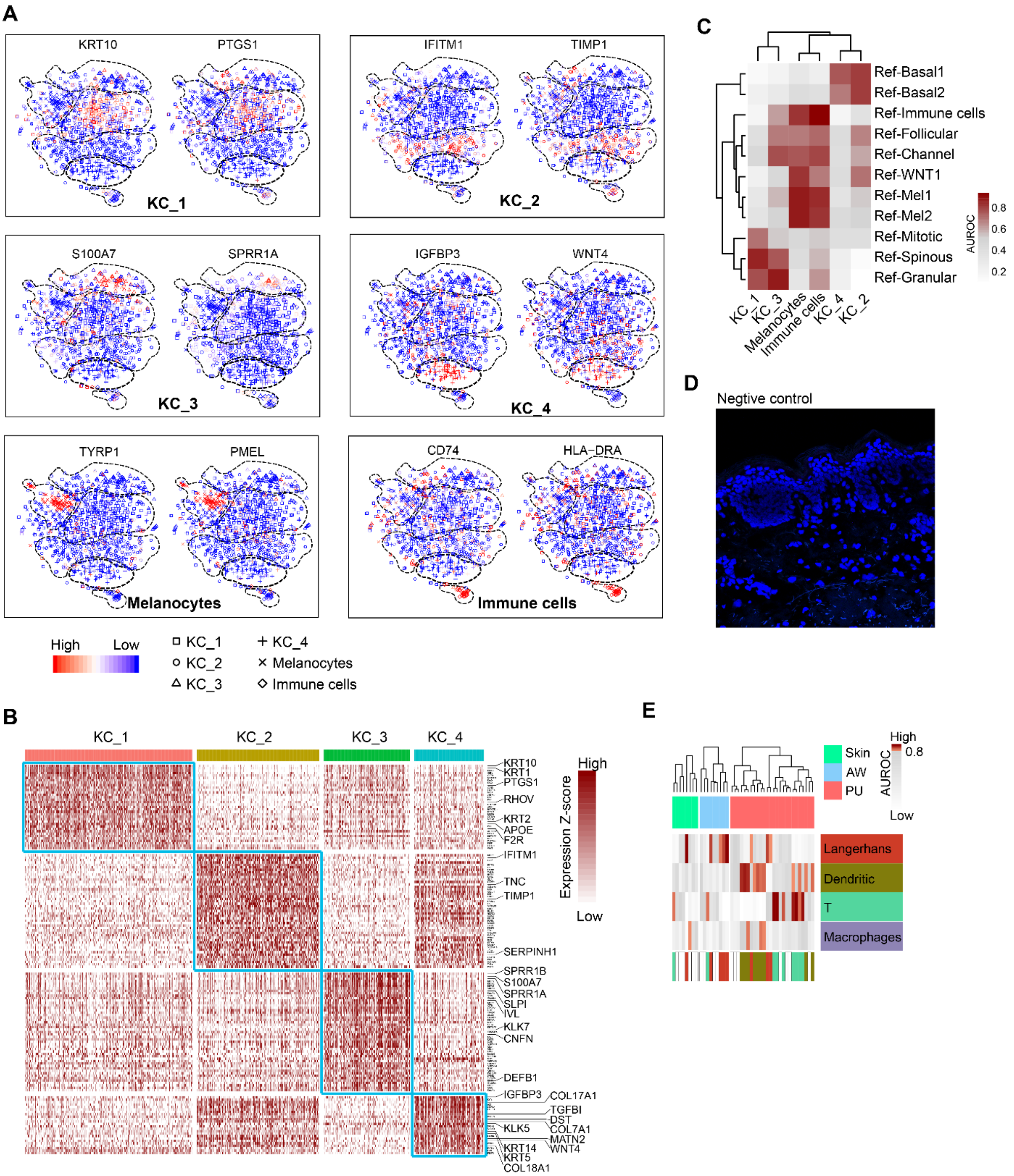


**Figure S3. Dissecting of the cellular composition of human skin and wound-edge epidermis by scRNA-seq. A**, tSNE projections of two selected maker genes for each cell cluster. **B**, Heatmap shows the expression of maker genes in four keratinocyte clusters. **C**, Comparison of our data with a published scRNA-seq dataset of human epidermis (Cheng, 2018). The similarity of two cell populations are indicated with a high receiver operator characteristic curve (AUROC) score. **D**, Negative control for IGFBP3 *in situ* hybridization staining (related to Figure 1D). **E,** Comparison of our scRNA-seq data of 43 immune cells with the published scRNA-seq dataset of 236 human epidermal immune cells (Cheng, 2018) identified three hematopoietic subsets in the skin and wound samples, i.e., Langerhans cells, myeloid dendritic cells, and αβT cells, with AUROC score > 0.8.

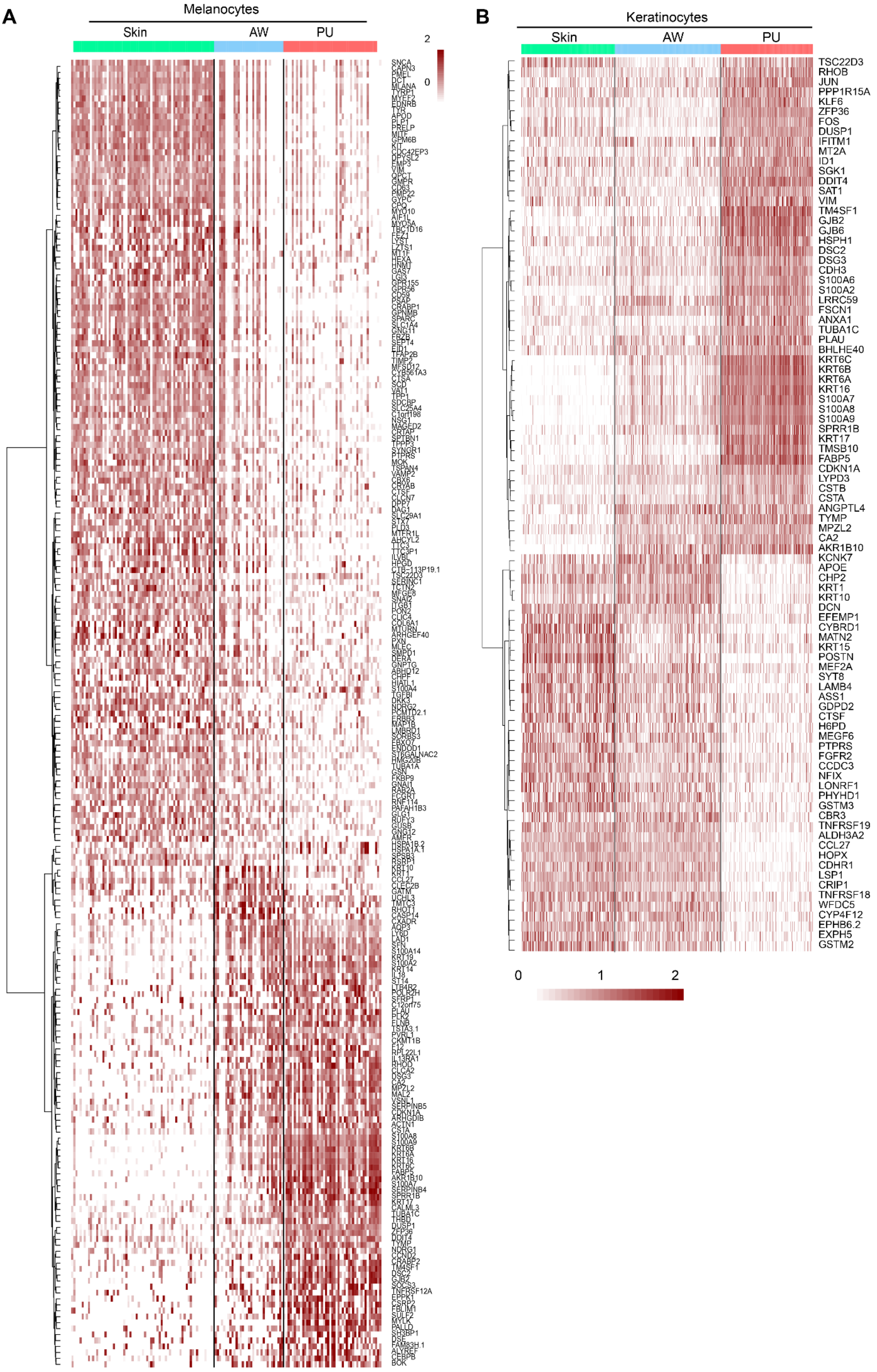
A
Melanocytes
Skin
AW
PU
2
0
B
Keratinocytes
Skin
AW
PU
TSC22D3
RHOB
JUN
PPP1R15A
KLF6
ZFP36
FOS
DUSP1
IFITM1
MT2A
ID1
SGK1
DDIT4
SAT1
VIM
TM4SF1
GJB2
GJB6
HSPH1
DSC2
DSG3
CDH3
S100A6
S100A2
LRRC59
FSCN1
ANXA1
TUBA1C
PLAU
BHLHE40
KRT6C
KRT6B
KRT6A
KRT16
S100A7
S100A8
S100A9
SPRR1B
KRT17
TMSB10
FABP5
CDKN1A
LYPD3
CSTB
CSTA
ANGPTL4
TYMP
MPZL2
CA2
AKR1B10
KCNK7
APOE
CHP2
KRT1
KRT10
DCN
EFEMP1
CYBRD1
MATN2
KRT15
POSTN
MEF2A
SYT8
LAMB4
ASS1
GDPD2
CTSF
H6PD
MEGF6
PTPRS
FGFR2
CCDC3
NFIX
LONRF1
PHYHD1
GSTM3
CBR3
TNFRSF19
ALDH3A2
CCL27
HOPX
CDHR1
LSP1
CRIP1
TNFRSF18
WFDC5
CYP4F12
EPHB6.2
EXPH5
GSTM2
0
1
2

**Figure S4. Differential gene expression in epidermal cells of acute wounds and pressure ulcers.** Heatmap illustrates the genes differentially expressed (fold change ≥ 2, adjust p value < 0.05) in melanocytes (**A**) and keratinocytes (**B**) from the skin, acute wounds (AW), and pressure ulcers (PU) at individual cell level. **C**, Immunofluorescence staining of cleaved caspase 3 (green) in the skin (n = 2), AW (n = 2), and PU (n = 5). Nuclei were co-stained with DAPI. Scale bar = 50 µm.

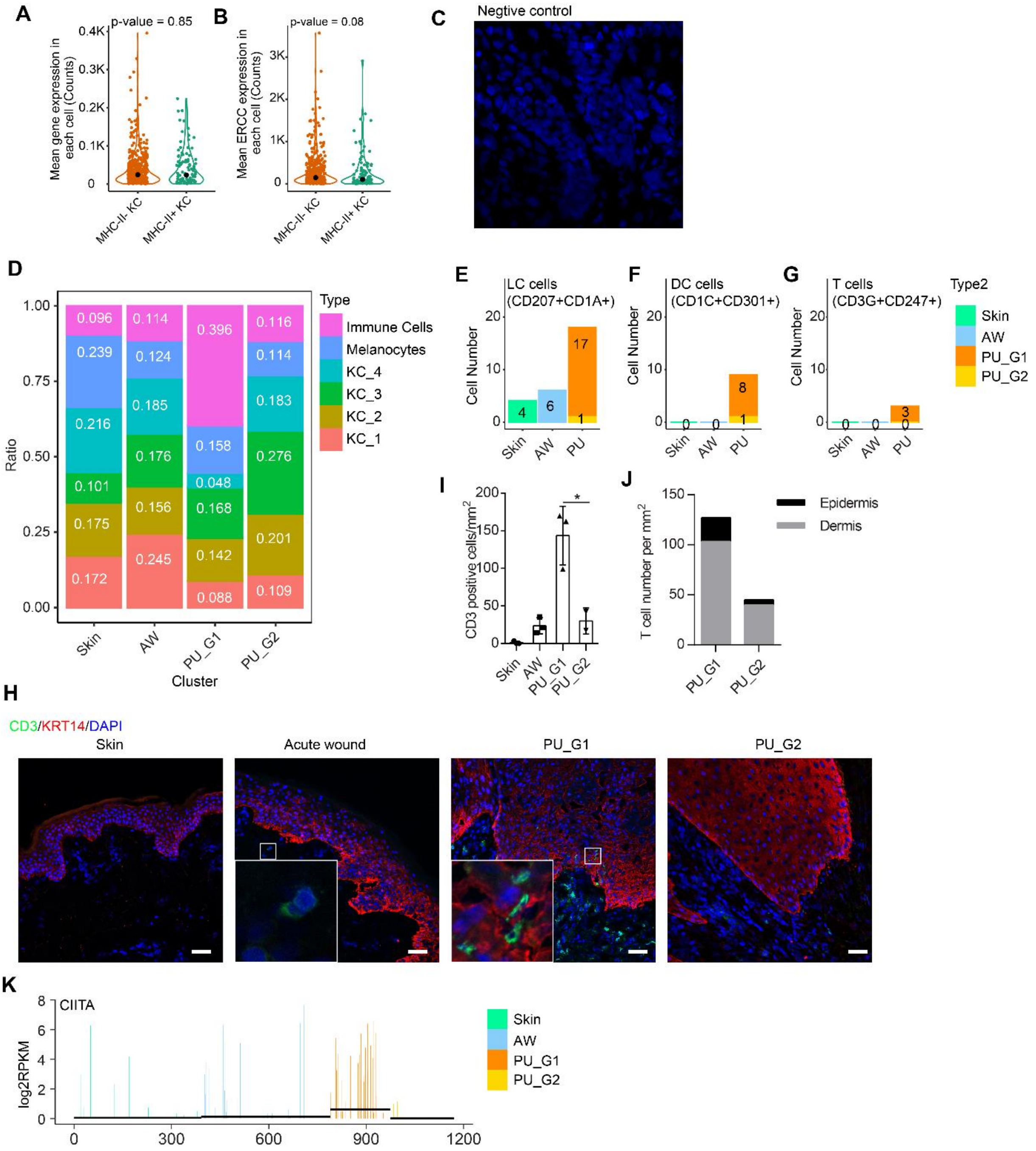


**Figure S5. Stratification of PU into two subtypes with distinct molecular features correlating with clinical outcomes.** Comparison of the mean gene expression (**A**) and the mean levels of ERCC spike-in RNA (**B**) between $MHCII^-$ and $MHCII^+$ keratinocytes. Wilcoxon test. **C**, Negative control for HLADR *in situ* hybridization staining (related to Figure 5E). **D**, Frequency distribution of the six epidermal cell clusters in the skin, AW, PU_G1 and PU_G2. **E-G**, The number of immune cells identified by scRNA-seq, including CD207+CD1A+ Langerhans cells (LC), CD1C+CD301A+ myeloid dendritic cells (DC), and CD3+ αβ T cells, in the skin, AW, PU_G1 and PU_G2 samples. **H**, Immunofluorescence (IF) co-staining of CD3 (green) and KRT14 (red) in the skin (n = 3), AW (n = 3), and PU (n = 5). Nuclei were co-stained with DAPI. Scale bar = 50 μm. **I-J**, The number of CD3 positive cells in epidermis and dermis were counted and normalized with the area of each field. **K**, The abundance of CIITA in the skin, AW, PU_G1 and PU_G2 is shown in all the cells. * $p<0.05$; Student's t-test (**I**).

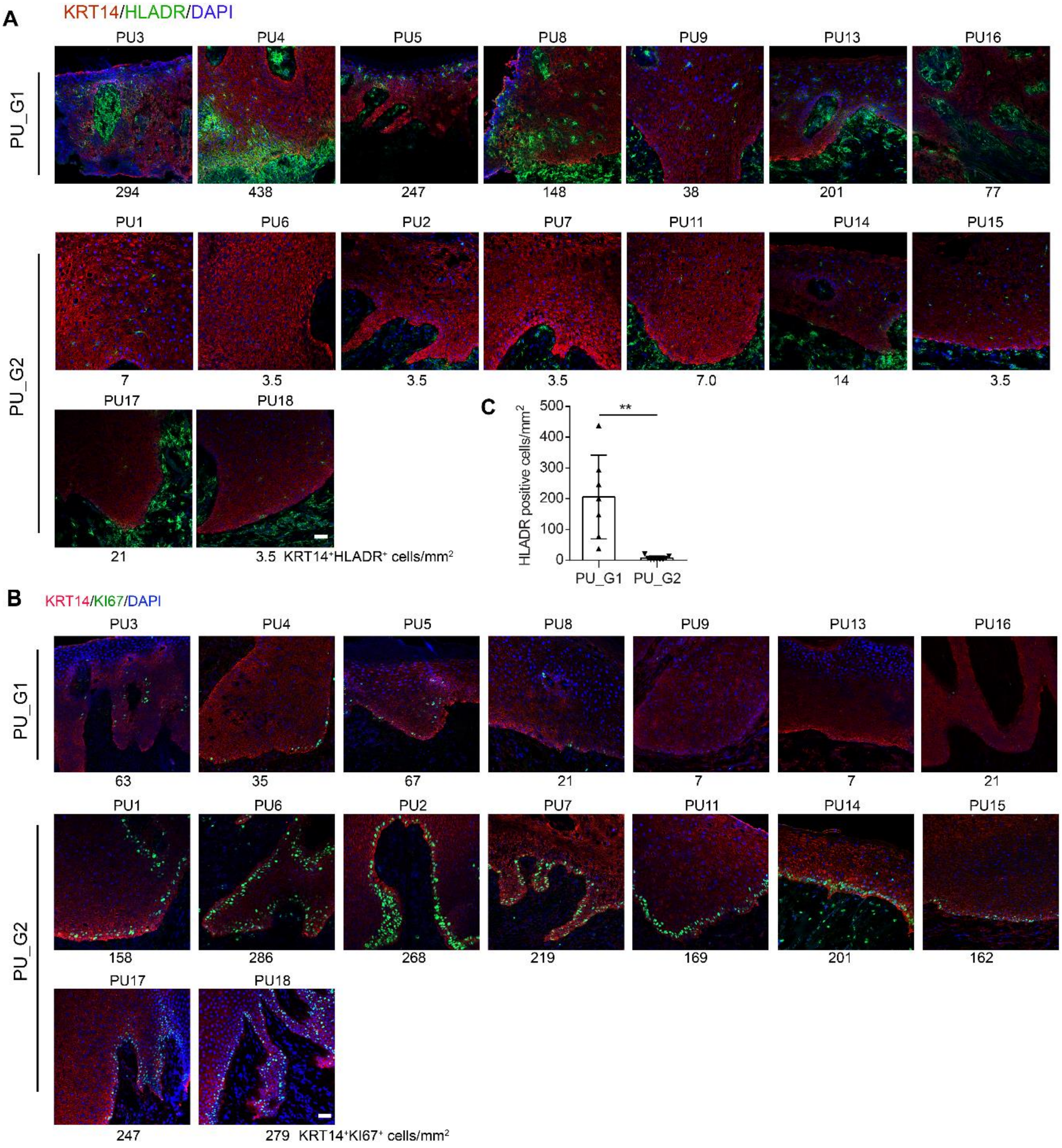


**Figure S6. Stratification of PU into two subtypes with distinct molecular features.** Immunofluorescence (IF) co-staining of HLADR (green) and KRT14 (red) (**A**) or KI67 (green) and KRT14 (red) (**B**) in pressure ulcer (PU) samples. Nuclei were stained with DAPI. Scale bar = 50µm. Based on the counts of positive stained cells in the epidermis (listed under each photograph), the PU samples were stratified into two subtypes. **C**, the number of HLADR positive cells in epidermis in PU_G1 and PU_G2 were counted and normalized with the area of each field. * $p<0.05$; Student's t-test (**C**).

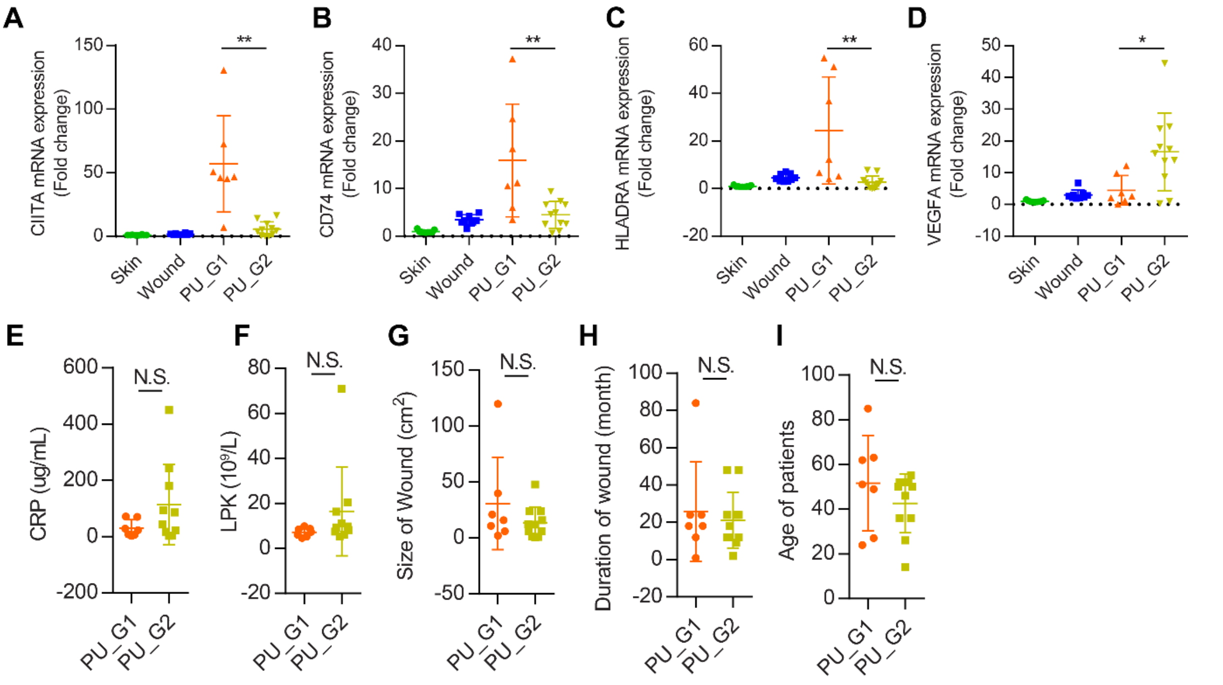


**Figure S7. Characterization of PU_G1 and PU_G2 samples. A-D**, QRT-PCR analysis of CIITA (**A**), CD74 (**B**), HLADRA (**C**) and VEGFA (D) expression in human Skin (n = 10), acute wound (n = 10), PU_G1(n = 7) and PU_G2 samples (n = 10). Comparison of the levels of circulating C-reaction protein (CRP) (**E**), leukocytes (LPK) (**F**), size of wound (**G**), duration of wound (**H**) and age (**I**) in two PU groups. * $p<0.05$, ** $p<0.01$, N.S. not significant, student t-test.

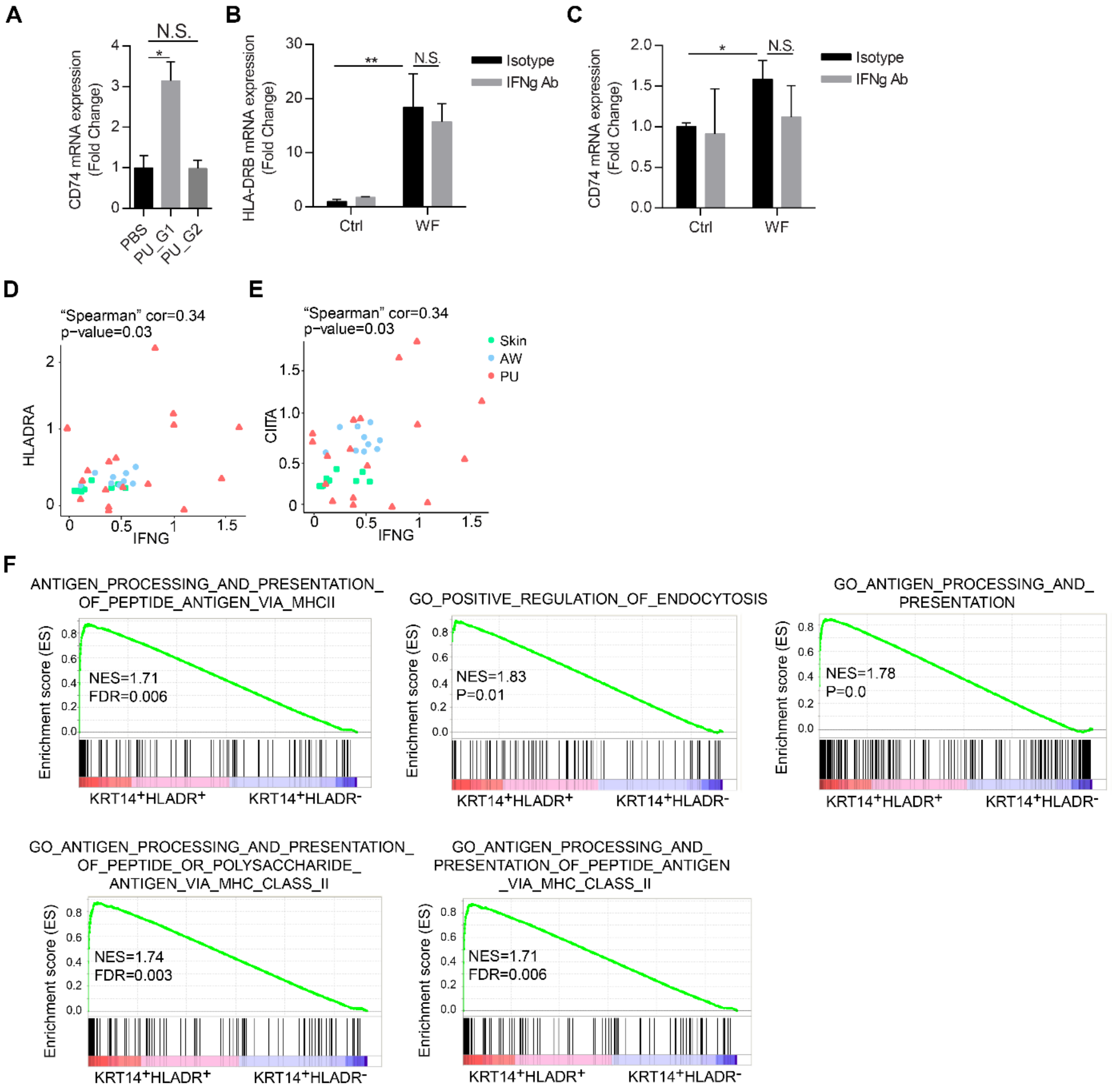


**Figure S8. IFNγ in PU wound fluids induces MHCII expression in keratinocytes.** **A**, qRT-PCR analysis of CD74 expression in human adult primary keratinocytes (HEKa) treated with wound fluid from PU_G1 and PU_G2 patients. **B**-**C**, human adult primary keratinocytes (HEKa) were treated with IFNg antibody 24 hours after treatment with wound fluid, the expression of HLA-DRB and CD74 was analyzed by QRT-PCR. **D**-**E**, Spearman's correlation analysis between IFNG and HLADRA (**D**) or CIITA (**E**) expression detected by qRT-PCR in the full-depth biopsies of human skin (n = 10), acute wounds (AW, n = 10), and pressure ulcers (PU, n = 17). **F**, GSEA of the genes related to MHC-II-mediated antigen presentation among the genes differentially expressed between CD74$^{high}$HLA-DRB$^{high}$ and CD74$^{-}$HLA-DRB$^{-}$ keratinocytes. * p<0.05, ** p<0.01, N.S. not significant, student t-test.

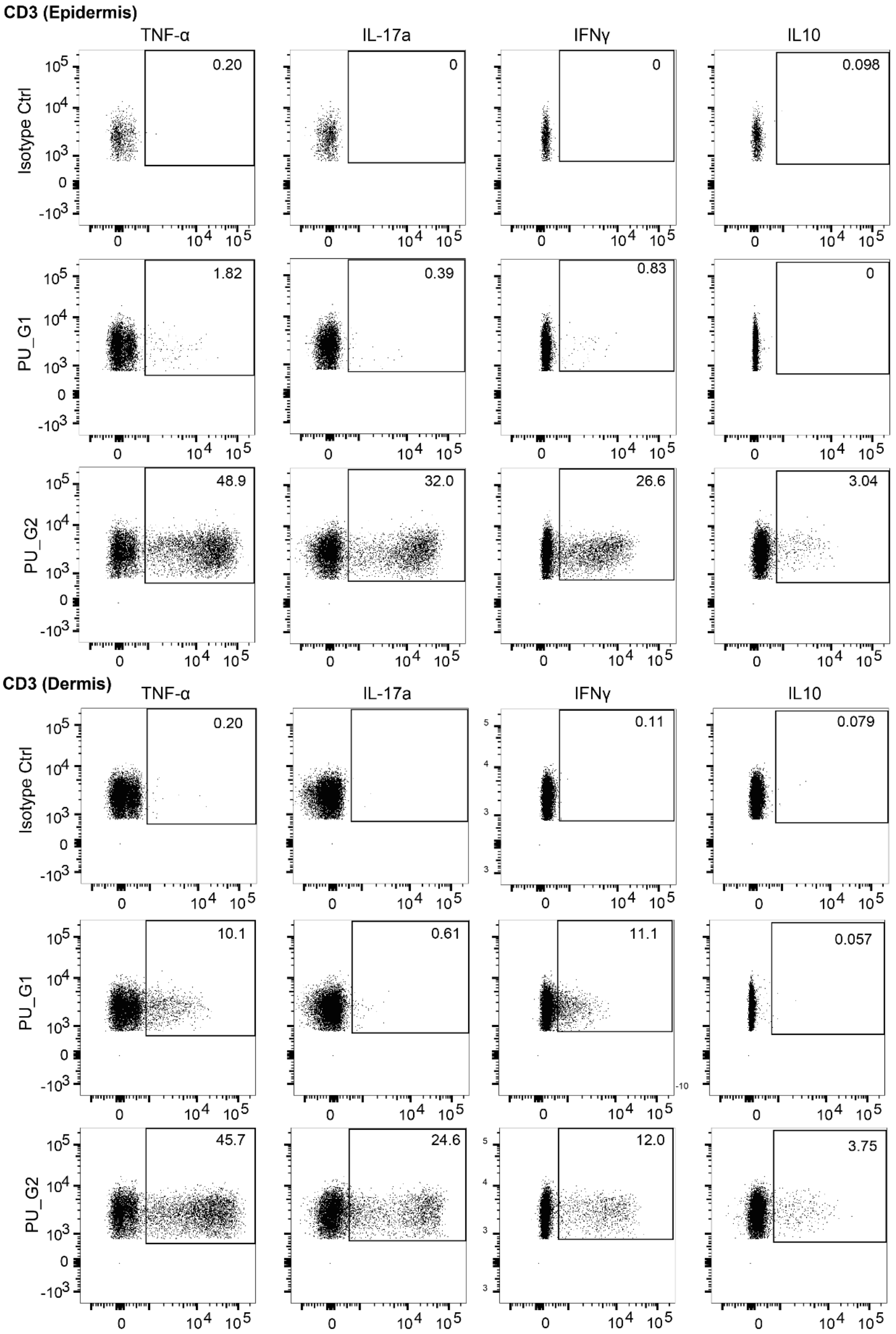


**Figure S9**. Flow cytometry analysis of TNFα, IL17a, IFNγ, and IL10 production in $CD3^{+}$ T cells from the epidermis (up) and dermis (down) of wound-edge biopsies of two PU groups.

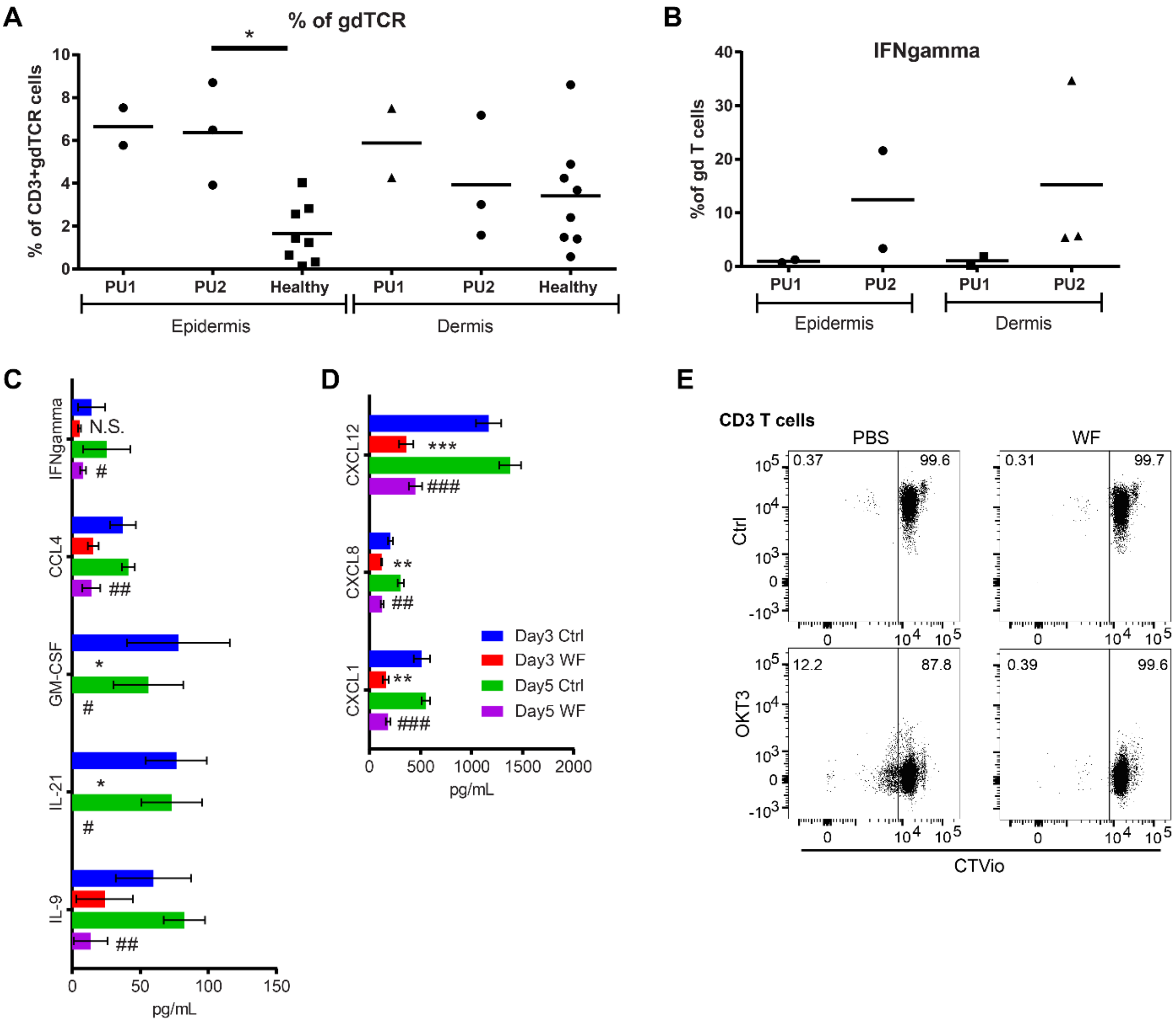


**Figure S10**. **Keratinocytes may function as an atypical antigen presenting cells and exert an inhibitory effect on T cell activation in PU wound-edges. A**, Flow cytometry analysis of percentage of gd T cells from the epidermis and dermis of the healthy skin (n = 8), PU_G1 (n = 2), and PU_G2 (n = 3) samples. **B**, Flow cytometry analysis of IFNgamma production in gd T cells from the epidermis and dermis of biopsies of PU_G1 (n = 2) and PU_G2 (n = 3) samples. **C-D**, ProcartaPlex immunoassay of cytokines in conditioned medium of autologous keratinocyte-T cell coculture for 3 and 5 days. Keratinocytes were pre-treated with or without wound fluid (WF). * Day 3 WF compared with Day 3 Ctrl, # Day 5 WF compared with Day 5 Ctrl. ## $p < 0.01$, ### $p < 0.001$, ** $p < 0.01$, *** $p < 0.001$, Student's t-test. **E**, Keratinocytes were pretreated with WF and co-cultured with autologous T cells activated with OKT3 (n=3 donors). T cell proliferation was analyzed by flow cytometry five days later.

#### Caption for movie S1

**Movie S1. RNAscope image of HLADR KRT14 double positive cells.** In situ hybridization of IGFBP3 (green) and KRT14 (red) mRNAs in chronic wounds. Nuclei were co-stained with DAPI. Three-dimensional videos were obtained from z-stacks after processing with Imaris image analysis software.
